# DeST-OT: Alignment of Spatiotemporal Transcriptomics Data

**DOI:** 10.1101/2024.03.05.583575

**Authors:** Peter Halmos, Xinhao Liu, Julian Gold, Feng Chen, Li Ding, Benjamin J. Raphael

**Author notes:** These authors contributed equally to this work, and the order is decided by a coin flip (Peter Halmos, Xinhao Liu).

## Abstract

Spatially resolved transcriptomics (SRT) measures mRNA transcripts at thousands of locations within a tissue slice, revealing spatial variations in gene expression and distribution of cell types. In recent studies, SRT has been applied to tissue slices from multiple timepoints during the development of an organism. Alignment of this *spatiotemporal* transcriptomics data can provide insights into the gene expression programs governing the growth and differentiation of cells over space and time. We introduce `DeST-OT` (**De**velopmental **S**patio**T**emporal **O**ptimal **T**ransport), a method to align SRT slices from pairs of developmental timepoints using the framework of optimal transport (OT). `DeST-OT` uses *semi-relaxed* optimal transport to precisely model cellular growth, death, and differentiation processes that are not well-modeled by existing alignment methods. We demonstrate the advantage of `DeST-OT` on simulated slices. We further introduce two metrics to quantify the plausibility of a spatiotemporal alignment: a *growth distortion metric* which quantifies the discrepancy between the inferred and the true cell type growth rates, and a *migration metric* which quantifies the distance traveled between ancestor and descendant cells. `DeST-OT` outperforms existing methods on these metrics in the alignment of spatiotemporal transcriptomics data from the development of axolotl brain.

**Code availability:** Software is available at https://github.com/raphael-group/DeST_OT

## 1 Introduction

Spatially Resolved Transcriptomics (SRT) technologies [37, 33, 28] measure gene expression simultaneously from thousands of cells or spots from a tissue slice, linking the gene expression measurement to the physical location within the tissue. These technologies enable exploration of tissue organization by analyzing cells within their native microenvironment, opening the door to the study of spatial biology [1, 17]. In some cases, SRT is applied to multiple slices from the same tissue. Joint analysis of multi-slice spatial data helps with the data sparsity problem in individual slices, enabling downstream analyses such as 3D differential expression or 3D cell-cell communication [23]. Multiple methods have been developed for alignment of multi-slice SRT data. For example, `PASTE` [45] integrates multiple slices from the same tissue and reconstructs the tissue gene expression in 3D, and `PASTE2` [24] extends `PASTE` to partially overlapping slices. `STalign` [8] is an image registration method finding a diffeomorphism between the H&E images of two spatial slices. `GPSA` [18] uses Gaussian processes to register spatial slices onto a common coordinate system, while `SLAT` [44] relies on graph neural networks and adversarial learning.

Another recent exciting application is to apply SRT to tissues taken from multiple timepoints of a developmental process [5]. Alignment of slices from multiple timepoints can provide insights into the gene expression programs governing the growth and differentiation of cells over space and time. However, alignment of spatiotemporal transcriptomics data presents unique challenges as there is a complicated interplay between proliferation and apoptotic cell dynamics in the sculpting of developing tissue [42]. Regions of the tissue may grow or shrink, creating many-to-one relationships between the spots from consecutive timepoints. Cells also change in gene expression and differentiate into new cell types during development. Moreover, slices no longer come from the same batch (individual or time-point) and thus may exhibit batch effects.

The existing methods for temporal alignment of single-cell data or for spatiotemporal alignment suffer from important limitations. `Waddington-OT` [34] aligns temporal single cell data for reprogramming datasets, but does not take spatial information into account. A recent preprint [21] describes `moscot`, a method that relaxes the OT formulation in `PASTE` to use *unbalanced optimal transport* [36]. `moscot` allows for cell growth and death using a curated set of proliferation and apoptosis genes as prior knowledge but optimizes an objective function that encourages static shape-matching. Other works [18, 44] do not necessarily quantify growth, and often have methodological and robustness limitations.

We introduce `DeST-OT`, a method to align spatiotemporal transcriptomics data that consists of SRT slices from multiple timepoints. `DeST-OT` proposes a novel *semi-relaxed* optimal transport framework, leading to unsupervised discovery of cell growth and apoptosis without relying on existing gene annotations. `DeST-OT` aligns differentiating cells along a manifold jointly defined by transcriptomic information and spatial information, leading to both biologically and physically valid alignments. `DeST-OT` accounts for spatiotemporal scenarios by modeling three orders of interactions between cells in a developing tissue, bridging a gap in the standard Fused Gromov-Wasserstein (FGW) OT objective [38].

We demonstrate the advantages of `DeST-OT` on both simulated spatiotemporal data and spatiotemporal data from axolotl brain development. To evaluate the performance of different methods we introduce the *growth distortion metric* quantifying the accuracy of the inferred cell growth within a tissue across timepoints, and the *migration metric* quantifying the distance that cells migrate during development under an alignment. We show that `DeST-OT` produces alignments that are more growth-aware on simulated data. `DeST-OT` alignments are more biologically realistic in terms of the growth inferred and the distance cells migrate compared to other methods. `DeST-OT` infers biologically valid cell type transitions on a spatiotemporal dataset of axolotl brain providing insights into the growth dynamics of brain development.

## 2 Methods

### 2.1 Formulation

A spatially resolved transcriptomics slice is represented by a tuple = (**X, S**). **X** ∈ ℕ ^n×p^ is the transcript count matrix where each row **x**_r_ is the gene expression vector of the corresponding spot, *n* the number of spots and *p* the number of genes measured. **S** ∈ ℝ ^n×2^ is the spatial position matrix, where the rows **s**_r_ encode the (*x, y*) coordinates of each spot. Given slices *𝒮*_1_ = (**X**^(1)^, **S**^(1)^) and *𝒮*_2_ = (**X**^(2)^, **S**^(2)^), measured on the same set of genes at timepoints *t*_1_ and *t*_2_, the goal is to derive an alignment matrix 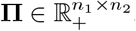, whose entry **Π**_ij_ is positive and gives the probability that the cell(s) in spot *i* of 𝒮_1_ are the progenitors of the cell(s) in spot *j* of *𝒮*_2_. The probabilities in the alignment matrix **Π**are derived to minimize some cost based on the gene expression at each spot and the spatial locations of aligned spots.

We use the mathematical tool of optimal transport (OT) to solve for **Π**. OT finds the most efficient way of moving mass between two distributions [30], and has previously been applied to single-cell alignment [11, 40] and spatial transcriptomics alignment [24, 45]. We seek to transport the mass of slice *𝒮*_1_ to *𝒮*_2_ where the mass is represented as a distribution over each slice’s spots. In the spatiotemporal setting, the amount of mass transferred between a spot at an earlier timepoint and a spot at a later one indicates how probable it is that the former spot is the ancestor of the latter. `PASTE` [45] uses OT to solve a related problem of static (non-temporal) spatial alignment, in which *𝒮*_1_ and *𝒮*_2_ are adjacent slices of the same tissue from the same timepoint, and minimizes the following objective function:

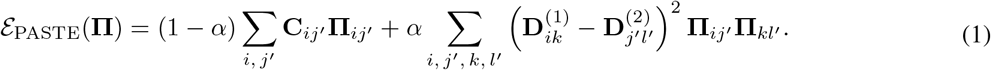

This objective function is a convex combination of two terms weighted by a balance parameter *α*. The first term, which we call the *feature term*, encourages matching spots with similar gene expression, and is also called the Wasserstein term in the OT literature [30]. We use the convention that a prime on an index, e.g. *j*^′^, refers to a spot in the second slice, while the absence of a prime denotes a spot in the first slice. The *ij*^′^-th entry of the matrix 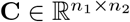 is the distance in expression space between expression vector **x**_i_ at *𝒮*_1_-spot *i* and expression vector 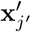 at *𝒮*_2_-spot *j*^′^. The second term, which we call the *spatial term*, encourages matching the intra-slice spatial distance between pairs of spots in each slice, and is called the Gromov-Wasserstein (GW) term [26, 31]. Matrices **D**^(1)^ and **D**^(2)^ are defined by intra-slice spatial distances **D**_ij_ = ∥**s**_i_−**s**_j_∥ _2_. The convex combination of the feature term and the spatial term is called the *Fused Gromov-Wasserstein* (FGW) objective [38]

`PASTE` optimizes (1) subject to the following constraints:

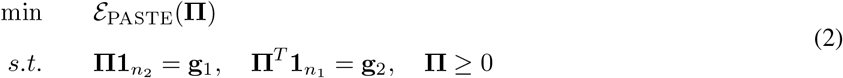

where 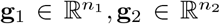 are uniform probability measures supported on the indices *i* ∈{1, …, *n*_1_} and *j*^′^∈ {1, …, *n*_2_} ; **1** is a vector of all one’s. These constraints are called *balanced* optimal transport (OT) (Fig. 1b) because the alignment matrix satisfies the marginals **g**_1_, **g**_2_ strictly.

**Figure 1:**
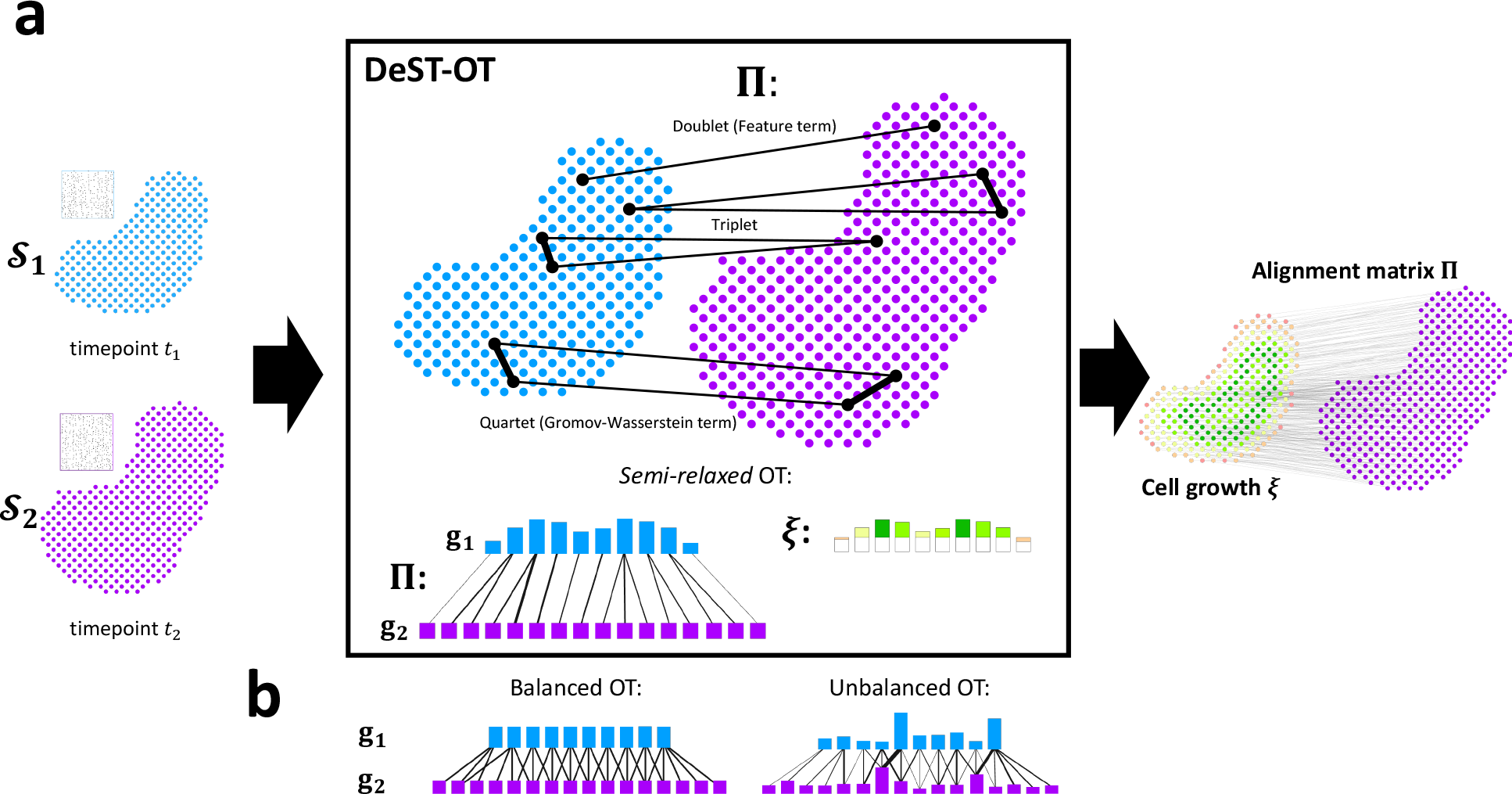
Overview of DeST-OT and semi-relaxed optimal transport (OT). (a) Given a pair of *𝒮*_1_ and *𝒮*_2_ from timepoints *t*_1_ and *t*_2_, respectively, `DeST-OT` infers an alignment matrix `Π`and a growth vector 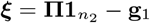 by solving a semi-relaxed optimal transport problem with doublet, triplet, quartet objective costs. Green entries in ***ξ*** indicate cell growth while red entries indicate death (b) Balanced OT, which fixes both marginals **g**_1_, **g**_2_, and unbalanced OT, which varies both marginals.

A recent preprint [21] introduced `moscot`, a modification of `PASTE` to use *unbalanced* OT, which is suggested to be helpful for spatiotemporal alignment. Specifically, `moscot` removes the equality constraints on the marginals, 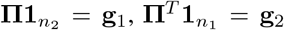 of (2), replacing these with two soft constraints in the form of Kullback-Leibler (KL) divergences:

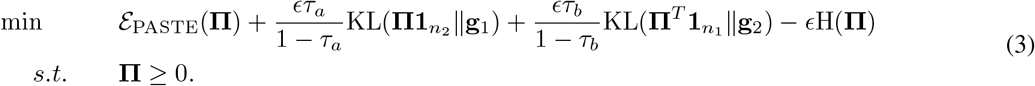

Here *τ*_a_, *τ*_b_ ∈ (0, 1) are hyperparameters determining the penalty for the marginals deviating from **g**_1_ and **g**_2_. H(·) is the entropy, where 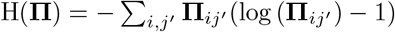. This entropic regularization accelerates the optimization of **Π**[9]. `moscot` has multiple limitations for spatiotemporal alignment. First, the method is supervised: to account for cell growth and death, `moscot` adjusts **g**_1_ over the first slice using the expression of a predefined set of marker genes, limiting its applicability to organisms and tissues with good prior knowledge. Secondly, the fully unbalanced formulation allows mass to shift around both marginals, limiting the interpretability of the growth information (Supplement § S1.4). Third, the spatial term is not amenable to tissue expansion because it prefers to align identical shapes: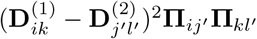 is minimized when the distances inside the square are the same.

`DeST-OT` uses *semi-relaxed* optimal transport (Fig. 1a) and optimizes a growth-aware objective function modeling different levels of interactions between cells, capturing the growth dynamics comprehensively. In the *semi-relaxed* optimal transport framework (Fig. 1), we relax only the constraint 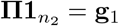 in (2), replacing it by a KL divergence in the objective function, while the other constraint 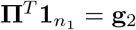 is kept. This ensures all of the spots in the second slice are mapped to from the first, while spots in the first slice can contribute a different amount of mass depending on whether they are growing or dying. We set both **g**_1_, **g**_2_ to assign equal weight 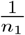 to each spot. That is, **g**_1_ is the uniform probability measure over spots in *𝒮*_1_, while **g**_2_ is a *positive* measure over spots in *𝒮*_2_. We define an interpretable *growth vector* ***ξ*** that represents a mass-flux across the two timepoints indicating the magnitude of growth and death for each spot in *𝒮*_1_,

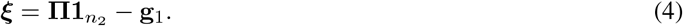

The growth vector 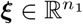 is the change in mass relative to a uniform prior **g**_1_ at each spot. The total sum of the entries of ***ξ*** is therefore 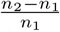, fixed in proportion to the total change of mass across slices. For a spot *i* at time *t*_1_, ***ξ***_i_ *>* 0 means that spot *i* has *>* 1 descendant in the second slice, and correspondingly, ***ξ***_i_ *<* 0 implies spot *i* has *<* 1 descendant in the second slice. The growth vector ***ξ*** is a change in mass over time – to convert this into a growth rate, one can take *J* = log(1 + *n*_1_***ξ***)*/*(*t*_2_ − *t*_1_) (Supplement § S2). See Supplement § S1.4 and § S3 for further discussion of the growth vector ***ξ***.

`DeST-OT` optimizes a growth-aware objective function subject to the semi-relaxed constraints. The objective cost of `DeST-OT` for finding an optimal spatiotemporal alignment matrix **Π**consists of three terms: a doublet term, a triplet term, and a quartet term. The doublet term, 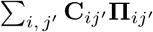, is the same as the feature term in `PASTE`, where 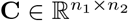 is an inter-slice gene expression distance matrix. The term compares the expression of two spots, one from each slice, hence we call it the *doublet* term of our objective.

The *quartet* term compares spot-pairs, with one pair from each slice. The quartet term is defined as 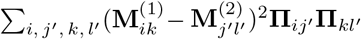, where each matrix **M**^(i)^ is defined to be the entrywise product of the square matrices **C**^(i)^ and **D**^(i)^ .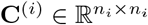is the distance in the expression space between each pair of spots on slice *i*. **D**^(i)^ is the intra-slice spatial distance matrix as in `PASTE`. The quartet term is equivalent to the spatial (GW) term of both `PASTE` and `moscot` but with matrices **M**^(i)^ that jointly model transcriptomic and spatial information, and with the semi-relaxed constraint applied to the gradient (Supplement § S4). We refer to **M** as *merged feature-spatial matrices*. While the spatial distance matrices (**D**^(1)^, **D**^(2)^) are appropriate for static alignment, they encode a rigid geometry that does not account for spatial deformations accompanying growth. On the contrary, `DeST-OT` matches a more flexible feature-smoothed geometry between the two slices, accounting for expansion or shrinkage of tissues during development.

The *triplet* term models the ancestor-descendant relationship between a single ancestor spot and multiple descendent spots, an essential relationship in growing tissues that is not well modeled by the other terms. When an ancestor spot differentiates into multiple descendant spots, these descendant spots should be close to each other in both physical space and feature space. Correspondingly, multiple ancestors should also be close. We make this notion precise by adding a *triplet* term to our objective: 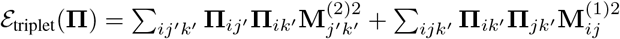. The entrywise squares on the **M** matrices match the form of the quartet term, upweighting the triplet summands which enforce the similarity of descendants and ancestors. Adding these terms to our objective function has a regularizing effect: **Π** is penalized for predicting distant descendants *j*^′^, *k*^′^ of the same spot *i* in the first slice, or for predicting distant ancestors *i, j* of the same spot *k*^′^ in the second slice. Distance is interpreted to be both spatial and transcriptomic due to the merged feature-spatial **M** matrices.

We call the sum *ε*^**M**^ of the triplet term and the quartet term the *merged feature-spatial* term, since both use merged feature-spatial matrices and encourage alignments to respect developmental dynamics in both physical space and gene expression space. Specifically, we define

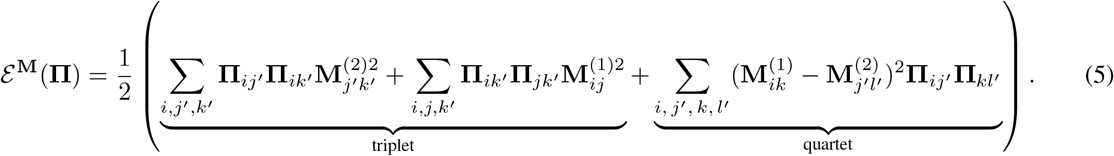

Combining all of the above, the `DeST-OT` objective function is:

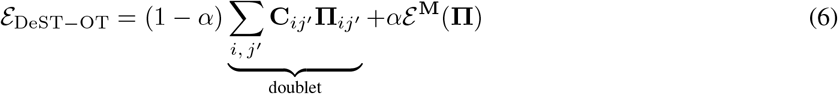

The combination of the doublet, triplet, and quartet terms captures lower to higher order of interactions between spots in a growing tissue (Fig. 1a). The `DeST-OT` optimization problem, with entropic regularization and the semi-relaxed constraints is

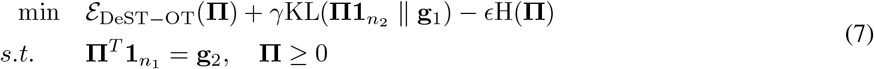

The balance parameter *α* balances the contribution of the feature term and the merged feature-spatial term to the alignment. *γ* governs the compliance of the semi-relaxed constraint, and *ϵ* governs the strength of entropic regularization. We discuss the effect of these hyperparameters in the Results section.

### 2.2 Optimization Using Sinkhorn

We solve the `DeST-OT` optimization problem by deriving a variant of the Sinkhorn algorithm, which has become the canonical way to compute OT alignments due to its speed [9]. One may convert an optimal transport problem with a general objective into the framework of Sinkhorn by adding an entropy regularization −*ϵ*H(**Π** ) to the objective function. Following the framework of Sinkhorn, the `DeST-OT` optimization problem (7) includes an entropy regularization term and we derive a set of updates from the KKT conditions for the semi-relaxed constraints to solve for **Π** . In practice, we take the dual of the semi-relaxed optimization to convert these updates into the log-domain [35], avoiding numerical overflow. The details of the optimization procedure are discussed in Supplement § S4.

### 2.3 Assessing alignment quality by cellular growth and migration

The true spatiotemporal alignment is often unknown, making it difficult to evaluate the accuracy of an alignment. We introduce two metrics to quantify the plausibility of an alignment: the *growth distortion metric* and the *migration metric*. The growth distortion quantifies the difference between the inferred growth and the proportional change of cell types in the two slices, given cell type labels for each spot in both slices. The migration metric quantifies how far cells “move” from the first timepoint to the second in a common coordinate framework describing the actual tissue. We say that an alignment **Π** is biologically valid when its growth distortion is ≈ 0, and physically valid if the migration distance is low.

#### 2.3.1 A metric of growth distortion

Given an alignment matrix **Π** , we define the *growth distortion metric* to quantify how well the growth vector ***ξ*** defined by equation (4) matches the observed change in proportion of given cell type labels over both slices. Formally, we are given a partition 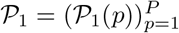 of spots in slice 1, where the set *𝒫*_1_(*p*) consists of all spots of cell type *p* at time *t*_1_, and a partition 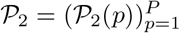 of spots at timepoint *t*_2_. The mass *m*_1_(*p*) of cell type *p* at time *t*_1_ is *m*_1_(*p*) = |*𝒫*_1_(*p*)|, the number of *t*_1_-spots with the label *p*. Likewise, the mass *m*_2_(*p*) of cell type *p* at time *t*_2_ is *m*_2_(*p*) = |P_2_(*p*)|. The change-in-mass for cell type *p* across these two timepoints is then *m*_2_(*p*) − *m*_1_(*p*). To ensure that the mass change has the same scale as the growth vector ***ξ***, we normalize the change-in-mass as 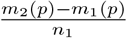 since `DeST-OT` marginals assign mass 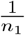 to each spot, while the counting measure used to define the *m*_t_(*p*) assigns mass 1 to each spot.

We define the growth distortion metric under two assumptions: first, that there are no cell type transitions between distinct cell types (we discuss how to relax this assumption shortly). Second, the burden of accomplishing the change in mass is shared equally across cells of the same type. This second assumption can be viewed as an entropy-maximizing assumption. Under these two assumptions, the “true” growth *γ*(*p*) at any *i* ∈ *𝒫*_1_(*p*) is

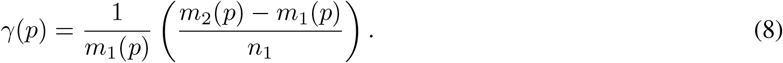

Note that summing these values over all *t*_1_-spots yields 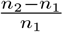, the total (normalized) change in mass across the two slices (Supplement § S5.1). The *growth distortion metric 𝒥*_*growth*_ of an alignment matrix **Π** with its associated ***ξ***, relative to cell type partitions *𝒫*_1_ and *𝒫*_2_, measures the total distortion between the inferred growth ***ξ*** and the true growth ***γ*** at each spot:

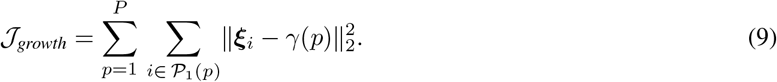

We generalize the growth distortion metric to the case when cell type transitions are present (but unknown) using a reverse-time transition matrix **T** ∈ ℝ ^P ×P^. This matrix acts on a vector of cell type masses, redistributing the mass **m**_2_ at time *t*_2_ to the ancestral cell types at *t*_1_ via the update **m**_1_ = **Tm**_2_. To compute the growth distortion metric for an alignment matrix **Π** , we use the following cell type transition matrix **T**:

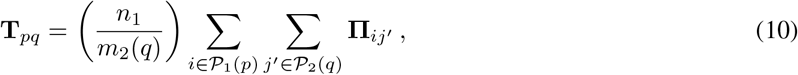

and prove in Proposition 2 of Supplement § S5.2 that the above **T** minimizes 𝒥_*growth*_ for a given **Π** across all **T**’s. That is, when we do not know the true cell type transitions, we compute the growth distortion of an alignment as the lowest distortion it could possibly achieve under any cell type transition.

#### 2.3.2 A metric of cell migration

We introduce a migration metric *𝒥*_*migration*_ of an alignment **Π** between two slices that quantifies the distance cells move under the alignment. This metric formalizes the intuition that the descendants of a cell tend to be close to their parent, particularly over short time intervals. Given an alignment **Π** and function *φ* : ℝ ^2^ → ℝ ^2^ that places slice *𝒮*_2_ intoa common-coordinate frame with slice *𝒮*_1_, we define the *migration metric* as:

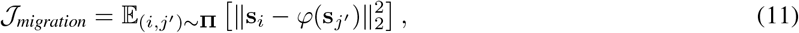

namely the average squared distance between spatial coordinate **s**_i_ in slice *𝒮*_1_ and transformed second-slice spatial coordinate *φ*(**s**_j_′) over pairs (*i, j*^′^) that are sampled proportionally to **Π** (column normalizing **Π** , as in Supplement Eq. (49)). In the results reported below, we use the function *φ*(**z**) = **Q**(**z** − **h**) for an orthogonal transformation **Q** and translation vector **h** ∈ ℝ ^2^ that solve a generalized Procrustes’ problem (Supplement § S6). This describes a rigid-body transformation relating the coordinate frames of slice *𝒮*_1_ and *𝒮*_2_.

## 3 Results

### 3.1 Evaluation on simulated ST data

We evaluated `DeST-OT` and `moscot` on simulated data from one- and two-dimensional tissue slices with eight-dimensional feature expressions for each spot. For each timepoint, the feature at each spot varies within each cell type. Details of the simulation of features are in Supplement § S7. Since `moscot` assumes the marginals used as input to its OT problem are already adapted to cell proliferation and apoptosis using prior knowledge, we set `moscot`’s marginal over *t*_1_ to account for changing cell type proportion in each experiment. `DeST-OT` uses semi-relaxed OT, and thus no prior knowledge of growth and death was required.

The first simulated dataset consisted of a pair of one-dimensional tissue slices, denoted as timepoint *t*_1_ and *t*_2_, each with 101 spots. There were two cell types across both slices. Slice *t*_1_ had 30 spots of cell type A and 71 spots of cell type B; slice *t*_2_ had 60 spots of cell type A and 41 spots of cell type B (Fig. 2ab). Therefore, cell type A grew by a factor of 2 from *t*_1_ to *t*_2_ and cell type B shrunk by roughly the same factor. We ran both `DeST-OT` and `moscot` on this pair of one-dimensional slices, varying the balance parameter *α* in the objective functions from *α* = 0.1 to *α* = 0.9, gradually placing more weight on each method’s spatial term (i.e. the merged feature-spatial term in `DeST-OT`, and the GW term in `moscot`) in the objective. We found that `DeST-OT` alignments are robust to varying *α*, always aligning cells to the correct cell types across timepoints (Fig. 2a). `DeST-OT` captures the true growth pattern of cells for all values of *α* because the merged feature-spatial term of `DeST-OT` incorporates both transcriptional and spatial information. On the other hand, `moscot` has greater difficulty capturing growing and shrinking cell types with larger *α*, as its spatial term emphasizes matching the shapes of the two slices as discussed in §2.1 (Fig. 2b). This demonstrates the effectiveness of `DeST-OT`’s spatiotemporal objective function, as well as the importance of `DeST-OT`’s semi-relaxed framework even when aligning slices with the same number of spots.

**Figure 2:**
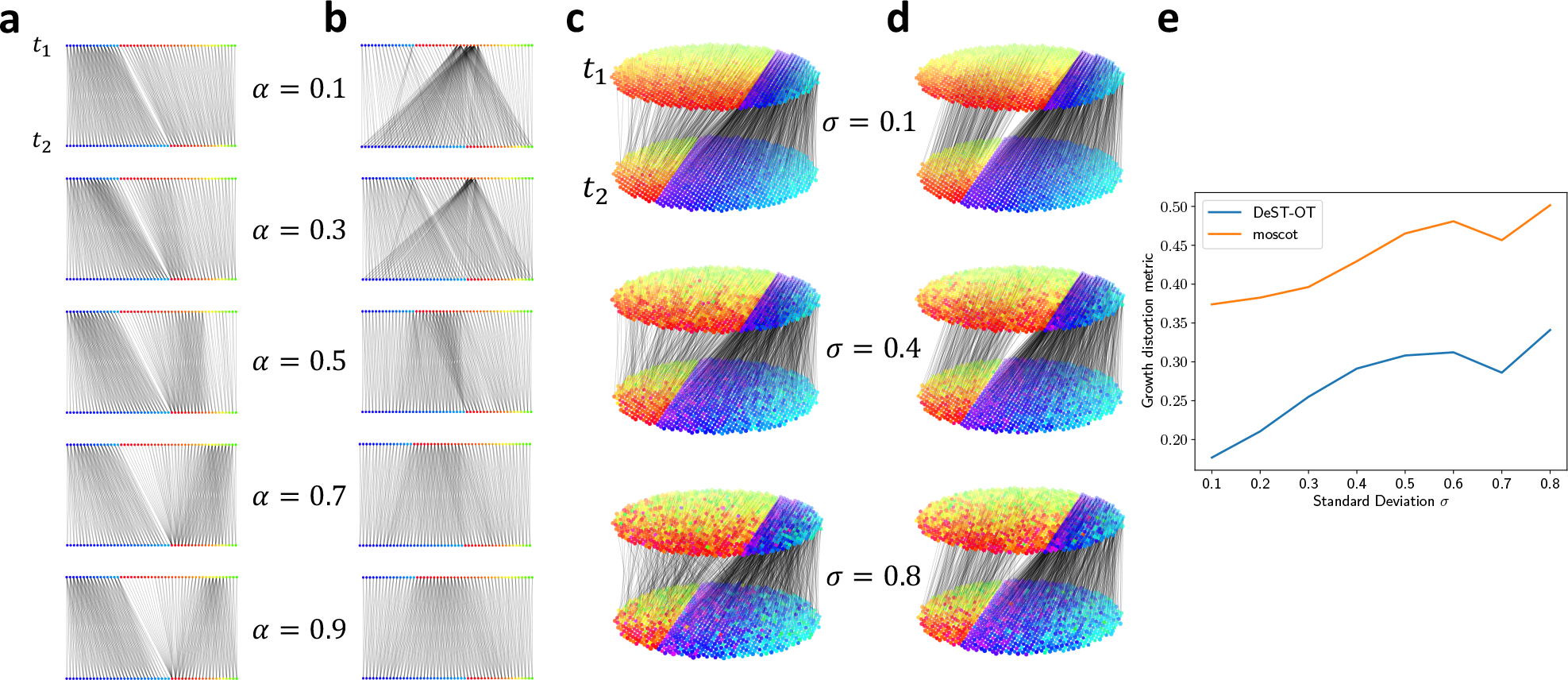
DeST-OT and moscot alignments on simulated data. **a**, `DeST-OT` and **b**, `moscot` alignment results for the balance parameter *α* ranging from 0.1 (mostly feature term) to 0.9 (mostly spatial terms), on 1D simulated slices. `DeST-OT`’s alignments indicate the cell type boundary, and color represents the polar angle made from two of the four non-zero coordinates in each cell type. **c**, `DeST-OT` and **d**, `moscot` alignments on 2D simulated slices with standard deviation of the expression noise *σ* = 0.1, 0.4, 0.8. Expression features are visualized similarly in 2D. **e**, The growth distortion metric for `DeST-OT` and `moscot` as a function of *σ*.

We next tested `DeST-OT` and `moscot` on a more realistic simulation with two-dimensional slices and feature expression noise. We generated two elliptical slices at *t*_1_ and *t*_2_ of the same size (988 spots), again with two cell types across the slices; cell type A occupies the right regions of the slices in (Fig. 2b), while cell type B is on the left. Slice *t*_1_ had 240 spots of cell type A and 748 spots of cell type B; slice *t*_2_ had 726 spots of cell type A and 262 spots of cell type B. Each cell type is characterized by a pair **f**_x,right_, **f**_x,left_ of eight-dimensional feature vectors for the *x*-direction, and another pair **f**_y,top_, **f**_y,bottom_ of feature vectors for the *y*-direction. The feature at a given spot in each cell type is a convex combination **f** (*x, y*) = *λ*_x_**f**_x,right_ + (1−*λ*_x_)**f**_x,left_ + *λ*_y_**f**_y,top_ + (1−*λ*_y_)**f**_y,bottom_ of the *x*-direction feature vectors the *y*-direction feature vectors. The coefficients *λ*_x_, *λ*_y_ are determined by the horizontal and vertical distance to the spot’s cell type boundary. This creates a consistent gradient of features within each cell type (Supplement § S7).

For (*x*_1_, *y*_1_) ∈ **S**^(1)^, (*x*_2_, *y*_2_) ∈ **S**^(2)^ we have that if 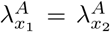 and 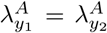 then **f**_A_(*x*_1_, *y*_1_) = **f**_A_(*x*_2_, *y*_2_) and the two spots should be aligned between timepoints 1 and 2 (likewise for cell type B). We then added zero-centered Gaussian noise with standard deviation *σ* independently to each feature dimension, with *σ* ranging from 0.1 to 0.8 in increments of 0.1. In addition to supplying `moscot` with the ground-truth adapted *t*_1_-marginal, we also set it to be fully unbalanced with *τ*_a_ = 0.99, *τ*_b_ = 0.999 as suggested by the tutorial for noisy data.

`DeST-OT` aligns ancestor cells to descendant cells correctly along the cell type feature gradients across timepoints (Fig. 2c), capturing cell growth and death. `DeST-OT` alignments are robust to noise as well, aligning cell types correctly even with added noise in spot features. While `moscot` produces straighter alignments (Fig. 2d) we found that it identifies the ancestors of cell type B at *t*_2_ to come from a small, similarly shaped region of cell type A at *t*_1_ even with perfect growth knowledge encoded in the marginal **g**_1_. Rather than aligning along a gradient of spatially expanding features, `moscot`’s spatial term prefers to align identical shapes, and is not suited to aligning tissue slices which grow and deform over time. `DeST-OT` consistently infers more accurate cell development as quantitatively shown by the growth distortion metric (Fig. 2e).

### 3.2 Axolotl brain development

We applied `DeST-OT` to analyze the developmental dynamics of the telencephalon, a region of the brain, in axolotl (*Ambystoma mexicanum*), a species of salamander. *Wei et. al*. [43] used Stereo-seq [5] to measure gene expression in the axolotl telencephalon at five development timepoints: three embryonic stages (44, 54, 57), Juvenile stage, and Adult stage (Fig. 3a). The slices grow in size, and progenitor cell types transition into mature cell types during the developmental process. We used `DeST-OT`, `moscot`, `PASTE`, `STalign`, and `SLAT` to infer an alignment between each pair of timepoints respectively, and computed the growth distortion and migration metric for each method (Fig. 4). Since new cell types appear at individual timepoints and the transitions between cell type during development are not annotated, we computed the growth distortion metric for each method under the cell type transition that minimizes their growth distortion as described in § 2.3. `DeST-OT` has the lowest growth distortion among all methods while maintaining low migration distances, demonstrating the quality of the `DeST-OT` alignments (Fig. 4). `moscot` and `STalign` achieve a low migration metric by shape-matching but have high growth distortions. `SLAT` has a low migration distance on this dataset as well, but cannot estimate cell growth and death accurately either.

**Figure 3:**
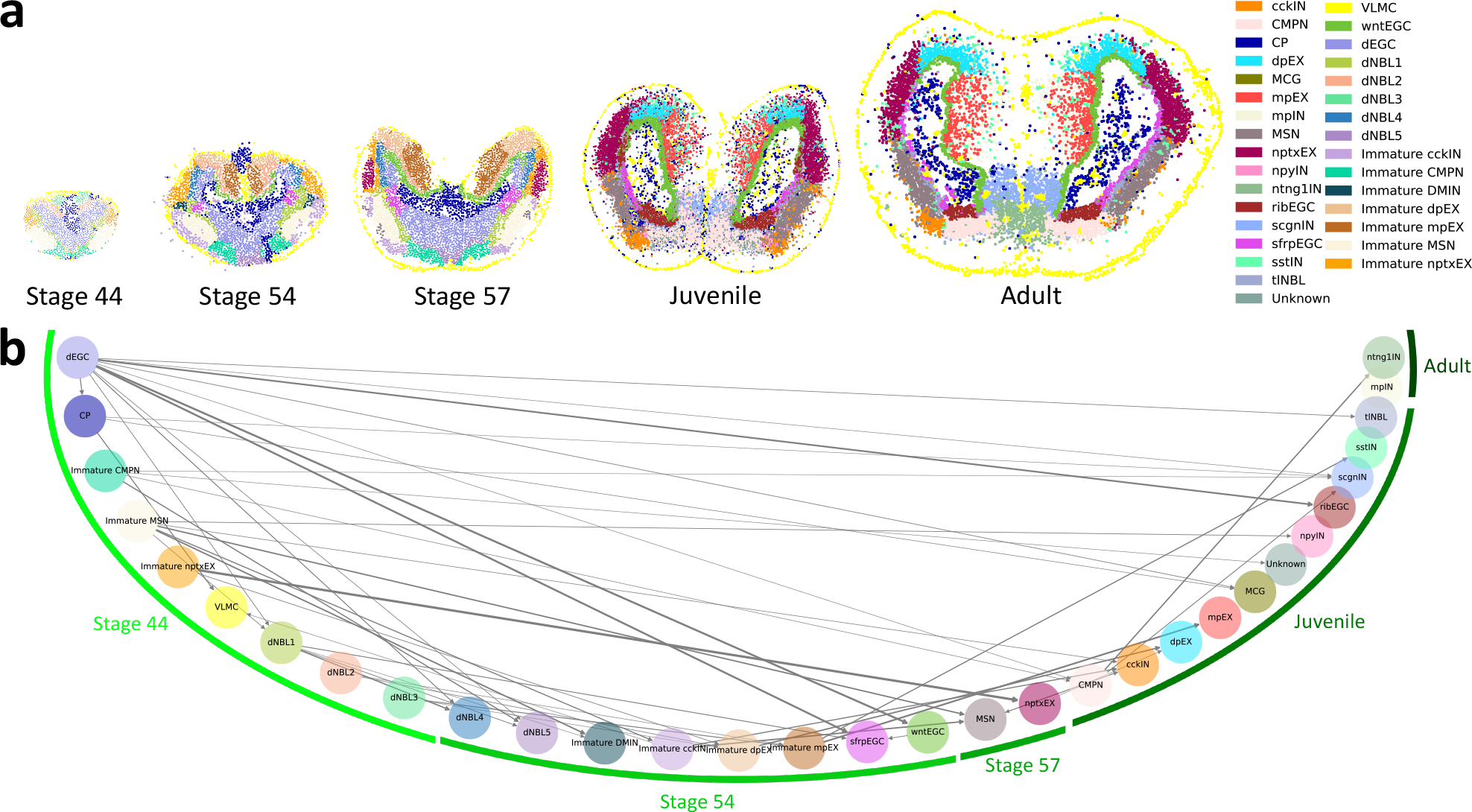
DeST-OT analysis of axolotl brain development. **a**, Stereo-seq data from axolotl brain sections at embryonic stage 44, embryonic stage 54, embryonic stage 57, Juvenile stage, and Adult stage, with cell type labels from [43] **b**, Cell type transition graph derived from `DeST-OT` alignments throughout axolotl brain development. The cell types are arranged in a half circle. A cell type is assigned to a developmental stage if it first appears in that stage. The width of the edges are proportional to the weight of transition. Self-loops are omitted.

**Figure 4:**
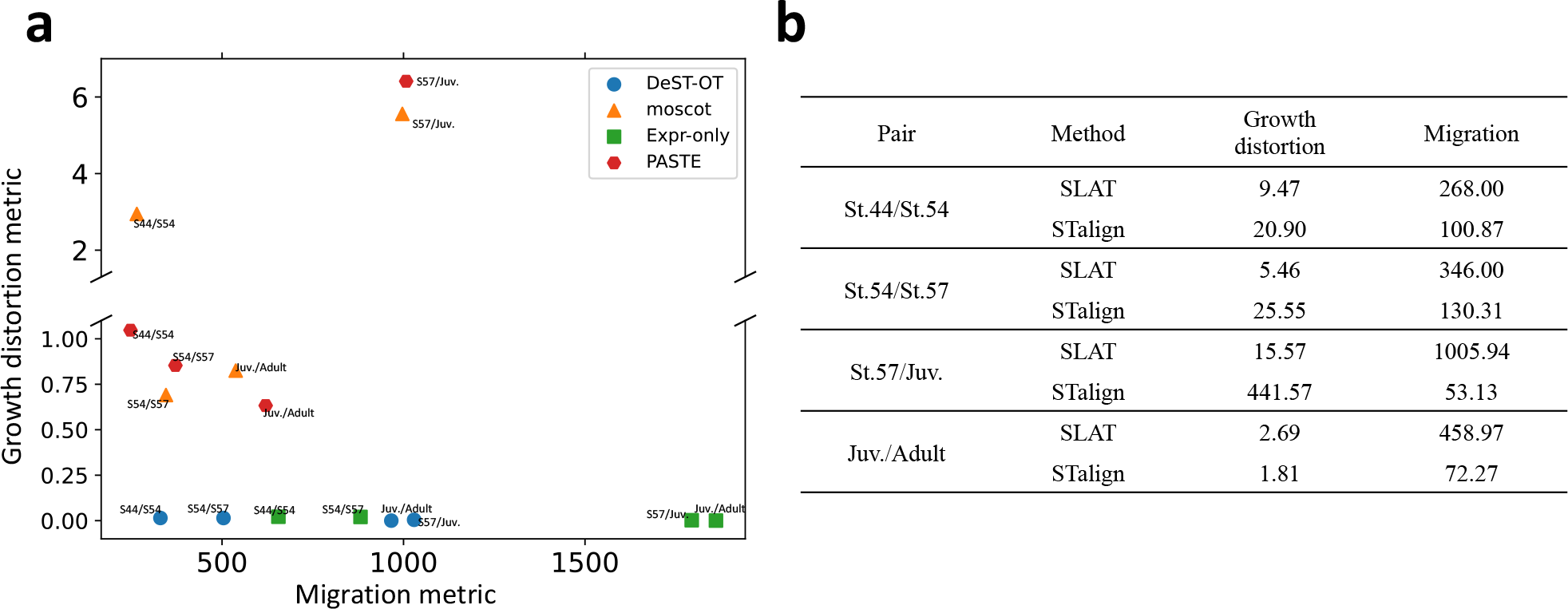
Alignment performance of various methods on the axolotl brain development dataset. **a**, The growth distortion metric and migration metric of `DeST-OT`, `moscot`, `Expr-only`, `PASTE` alignments on all pairs of axolotl brain development timepoints. **b**, The growth distortion and migration metric of `SLAT` and `STalign` alignments on all pairs of axolotl brain development timepoints. Not included in the plot in **a** because of high growth distortion.

We examined the cell type transition matrix **T** between each pair of adjacent timepoints derived from the `DeST-OT` alignment (§ 2.3) and the cell type annotations from [43] (Fig. S1). From these transition matrices, we derive a weighted directed graph showing all frequent transitions (*>*20%) (Fig. 3b). Many of the `DeST-OT` inferred cell type transitions are consistent with previously reported developmental trajectories. For example, we found that among all cell types, dEGCs (developmental ependymoglial cells) give rise to the largest number (11) of descendent cell types, consistent with previous studies which suggest that EGCs are equivalent to neural stem cells in mammals and contribute to neurogenesis during brain development [19, 20, 3]. Furthermore, immature cell types expressing early developmental markers disappear from the juvenile stage onward (Fig. 3a), and `DeST-OT` confirmed that immature cells of each cell type, such as CMPN or nptxEX, transition into their respective mature cell types. Previous studies suggested a potential lineage transition from EGCs to neuroblasts (NBLs) [29], and a transition from dEGC to NBL cell types that appear after stage 44 (dNBL4, dNBL5, tlNBL) is also found by `DeST-OT`. Finally, dEGCs disappear at stage 57, and `DeST-OT` predicts that it transitions mostly into ribEGCs located in the ventricular zone, which is the same spatial region where dEGCs located before and consistent with the findings in [43]. We observe a directed cycle between cckIN and MSN, probably because the two cell types are mixed together in the striatum region. There is no directed edge going into mpIN because it has multiple progenitor cell types and no cell type passes the threshold (20% in this case) for including an edge giving rise to mpIN (Fig. S1), indicating a diverse origin of mpIN.

We found that the growth patterns of individual cells inferred by `DeST-OT` are more biologically reasonable than those inferred by `moscot` (Fig. 5). For the three embryonic stages, `DeST-OT` infers that all cells are growing and identifies differential growth patterns of tissue regions: the outer part of the telencephalon, mostly occupied by immature cell types, grows faster than the inner part. In contrast, the growth patterns inferred by `moscot` tend to be sparser, with a few “representative” cells have a high growth and thus a large number of descendants in the next timepoint, while most other cells are dying, which is not realistic for embryonic tissues. This is a computational artifact of the fully unbalanced OT formulation which allows a few “best” cells from the two slices to be aligned, hence does not fit spatiotemporal data where all descendant spots should be aligned.

**Figure 5:**
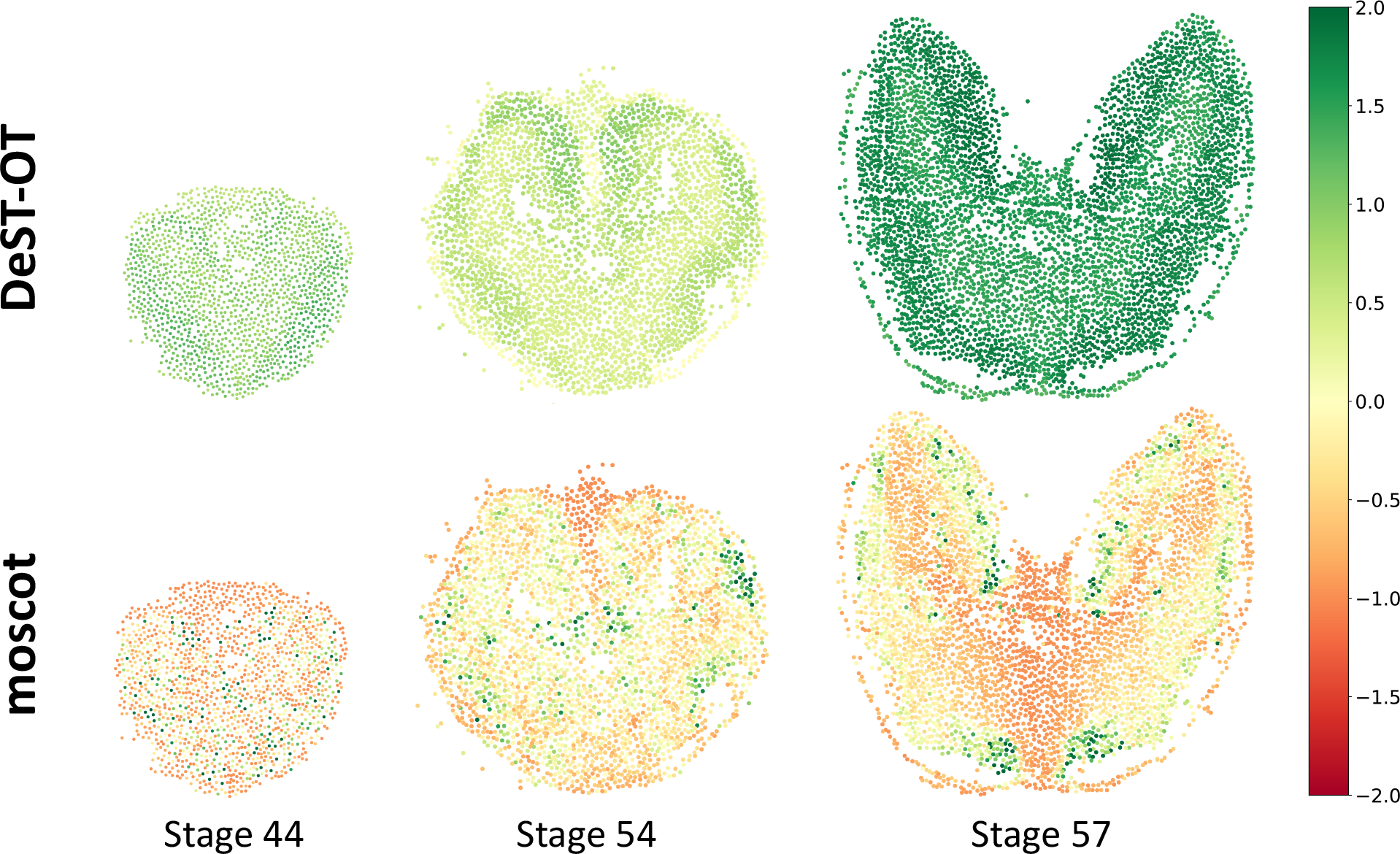
Growth patterns of axolotl brain. The growth of cells inferred by `DeST-OT` and `moscot` on the three embryonic stages of axolotl brain development. Growth vector ***ξ*** is normalized to unit of number of cells.

## 4 Discussion

We introduce `DeST-OT`, a method for aligning spatiotemporal transcriptomics data and for inferring cell proliferation and apoptosis. Using a semi-relaxed optimal transport framework and an objective cost designed for spatiotemporal data, `DeST-OT` finds an alignment between progenitor and descendent cells in developmental spatial transcriptomics data, infers growth and death rates, and infers cell type transitions during tissue development. To quanitfy the performance of `DeST-OT` and other spatiotemporal alignment methods, we introduce the migration metric to quantify the distance cells travel under a spatiotemporal alignment and the growth distortion metric to quantify how accurately an alignment infers growth relative to ground-truth cell type annotations. We show on simulated data that `DeST-OT` outperforms other methods and infers cell growth and death accurately. We use `DeST-OT` to study axolotl brain development and confirm previously reported lineage transitions. `DeST-OT` alignments can elucidate developmental dynamics and may lay the ground for the discovery of their molecular basis. We also demonstrate that `DeST-OT` alignments are more biologically and physically realistic than competing methods.

Future work includes the evaluation of `DeST-OT` on other spatiotemporal datasets. There are currently few such publicly available datasets, but analysis of another unpublished dataset is ongoing and will be included in a future revision. `DeST-OT` will be a useful tool for biologists to gain new insights in spatiotemporal processes such as development and reprogramming, discovering new temporally and spatially dependent biological phenomena.

## 5 Acknowledgements

This research was supported by NIH/NCI grant U24CA248453 to B.J.R. J.G. gratefully acknowledges support from the Schmidt DataX Fund at Princeton University made possible through a major gift from the Schmidt Futures Foundation.

## Supplement

### S1 Formulation

In vanilla optimal transport, and in the original formulation of `PASTE` [45], one has two constraints on alignment matrix **Π** , given by probability measures 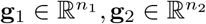 the row-sum of **Π** must be **g**_1_, and the column-sum of **Π** must be **g**_2_. Measures **g**_1_ and **g**_2_ are supported on indices *i* ∈ {1, …, *n*_1_} and *j*^′^ ∈ {1, …, *n*_2_}, enumerating the spots of two slices of spatial transcriptomics data, *𝒮*_1_ = (**X**^(1)^, **S**^(1)^) and S_1_ = (**X**^(2)^, **S**^(2)^). The rows 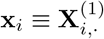 of **X**^(1)^ and 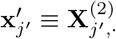 of **x**^(2)^ are indexed by spots, and each row is the expression vector at the given spot. Likewise, the rows 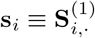 of **S**^(1)^ and 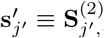 of **S**^(2)^ are the spatial coordinates of each spot. We use the convention that a prime on an index, e.g. *j*, refers to a spot in the second slice, while the absence of a prime on an index denotes a spot in the first slice.

#### S1.1 Context

In `PASTE`, a convex combination of standard inter-slice expression cost and a Gromov-Wasserstein cost are used (this combination referred to as *Fused Gromov-Wasserstein (FGW)*). The corresponding optimization problem is:

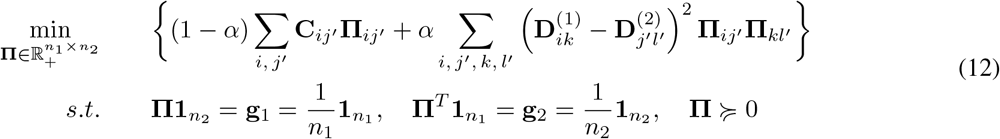

Above, 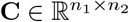 is the inter-slice (gene expression) feature-distance matrix 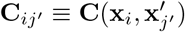 whose *ij*^′^-th entry is the distance in expression space between the expression vector **x**_i_ at spot *i* in the first slice and the expression vector 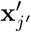 at spot *j*^′^ in the second slice. Matrices **D**^(1)^ and **D**^(2)^ in (12) are intra-slice (spatial) distance matrices. Define the 4-tensor 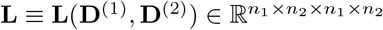 entry-wise via

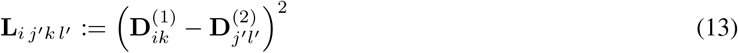

We let ⊗ denote the tensor-matrix multiplication operator, such that for the alignment matrix 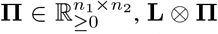 denotes the *n*_1_ × *n*_2_ matrix whose *ij*^′^-th entry is 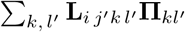. Let ⟨·,·⟩_*F*_ denote the Frobenius inner product of matrices. In this notation, the objective function (12) can be reformulated as follows:

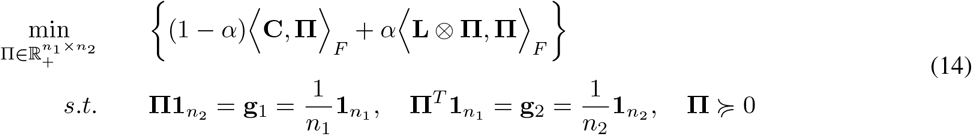

with **L** as in (13). The only difference between `PASTE2` [24] and the original `PASTE` is the inclusion of the constraint 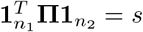, which goes along with a relaxation of the two marginal constraints to inequalities:

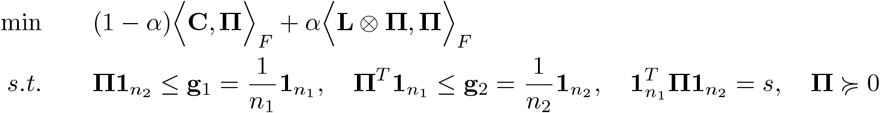

Note that by considering the dual problem, *partial optimal transport*, in which there is a constraint *s* ∈ (0, 1) on the total mass 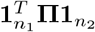 transported, can be seen as an instance of unbalanced optimal transport associated to the total variation *φ*-divergence (see [36] § 4.2). *Unbalanced optimal transport* refers to the version of (14) obtained by deleting the two hard constraints, 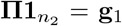 and 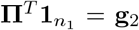, on the marginals of **Π** , and instead adding to the cost function in brackets a pair of “soft constraints” in the form of two terms: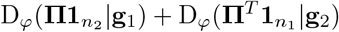.

Here, *φ* : (0, ∞) → [0, +∞] is a so-called *entropy function*, namely it is convex, positive, lower-semi-continuous, and with *φ*(1) = 0. Let D_*φ*_(·|·)denote the associated *φ*-divergence, defined for positive measures *α, β* on some common space *𝒳* as 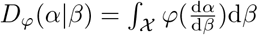, where we have supposed *α* and *β* are mutually absolutely continuous, for instance. When *φ*(*p*) = *p* log *p* − *p* + 1, one recovers the *Kullback-Leibler (KL) divergence*, which we denote by D_*φ*_(·|·) = KL(·∥·) in this case.

#### S1.2 DeST-OT objective

Our view of an OT objective function as a *Hamiltonian*, assigning an energy to a given transport plan **Π** , motivates the terminology we use for its components below. The first change we address in the formulation of `DeST-OT`, relative to `PASTE`, is the constraints on the optimization together with our calibration of mass on the second slice. Specifically, we relax the constraint 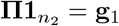, while preserving the other. This fixes the total mass transported by **Π** to be the total mass of the second (usually larger, in the spatiotemporal setting) slice, which is equipped with positive, *not necessarily probability*, measure 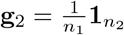. This allows the **Π** -marginal over the first slice, namely 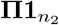, to be non-uniform,while fixing its total mass to the value 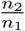. One is naturally lead to introduce a *mass-flux term*,

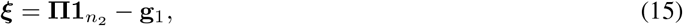

as this object becomes non-zero as soon as we relax the first marginal constraint. In section S1.4, we discuss the value of ***ξ***. The non-uniformity in 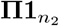 reflects that not all spots in the first slice are necessarily contributing the same amount to spots in the second slice – different cell types may have different growth rates.

The second change we address concerns the matrices used by the Gromov-Wasserstein (GW) term E_GW_(**Π** ) =

⟨**L**⊗**Π, Π**⟩. This term favors **Π** matrices which are nearly isometries between the two slices, as whenever 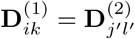, the corresponding summand of the GW term E_GW_(**Π** ) vanishes. For a more general pair of intra-slice cost maineqtrices, (symmetric, non-negative matrices) 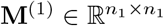 and 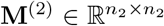, we generalize (13), defining the 4-tensor 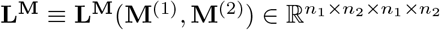 entry-wise via

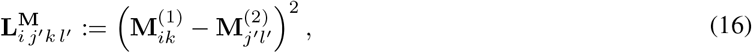

and we define the analogous GW energy (using **M**^(i)^ in place of the **D**^(i)^). We also refer to it as a *quartet* energy, as each summand defining it corresponds to two pairs points:

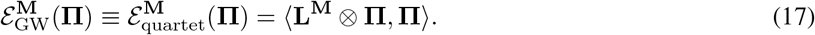

We define the matrices **M**^(i)^ from a pair of expression distance matrices for the first and second slice, 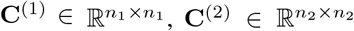. These intra-slice feature matrices **C**^(i)^ are distinct from their inter-slice counterpart 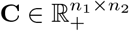 which contains the distance in (expression) feature space from spots in slice 1 and slice 2, and is therefore not a symmetric, square matrix in general. We specifically choose **M**^(i)^ to be the Hadamard product of matrices **C**^(i)^ and **D**^(i)^, so that entry-wise, one defines 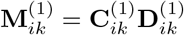 and 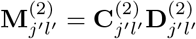. We refer to the **M**^(i)^ as *merged feature-spatial matrices*.

To 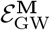, we add a symmetric pair of terms 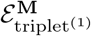 and 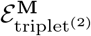 of the form

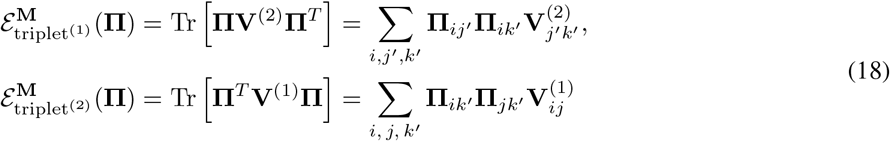

When **V**^(2)^ = (**M**^(2)^)^2^, i.e. entry-wise squaring of the merged feature-spatial matrix, which is why the superscript indicates the same dependence on the **M**-matrices as the GW term. Note that 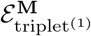 is identical to the terms in the sum defining 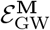 which have *i* = *k*, which is why we name it for the first slice instead using the index of the merged feature-spatial matrix involved in the definition. This energy 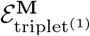 favors **Π** which predict a localized (in space, and in gene expression space) set of possible ancestors for each spot in the second slice.

Likewise, when **V**^(1)^ = (**M**^(1)^)^2^, energy 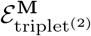 is identical to the terms in the sum defining 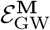 for which *j*^′^ = *l*^′^, and so is named for the second slice. Transport plans **Π** with low 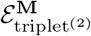 -energy are more likely to predict that descendants of a given spot in the first slice are close (spatially, and in feature-distance) in the second slice. Together, 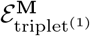 and 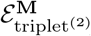 encourage spot-wise continuity in **Π** , without penalizing **Π** for spatial distortions intrinsic to tissue growth, or for change in expression characteristic of development. We refer to the sum of these terms as the *triplet* energy between triplets of points, ε_triplet_:

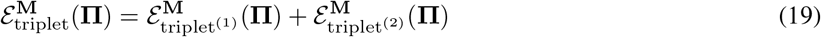

Combining these changes, and adding in a term of entropy regularization, the formulation ^1^ used by `DeST-OT` is:

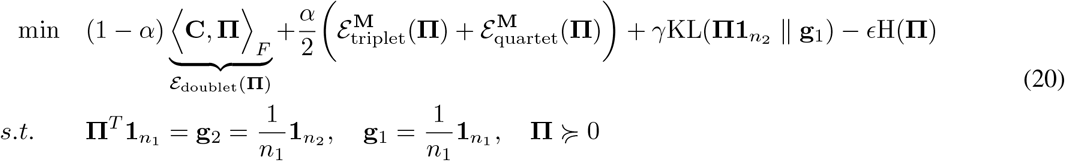

with **L**^**M**^ defined in (16), and the notation **L**^**M**^ ⊗ **Π** as defined just below (13). The choices of **V**^(1)^ = ((**M**^(1)^)^2^ and **V**^(2)^ = ((**M**^(2)^)^2^ as above enable us to write the sum of our triplet and quartet energies in (20) as

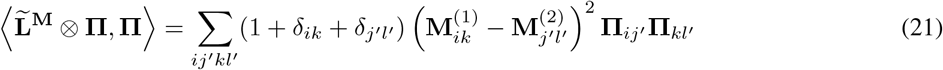

where *δ*_ik_ is 1 if and only if *i* = *k* and is 0 otherwise. The 4-tensor 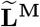 is defined implicitly by the above display, and the objective function used by `DeST-OT` can now expressed concisely as follows, though (20) is the expression we use in practice for computing gradients:

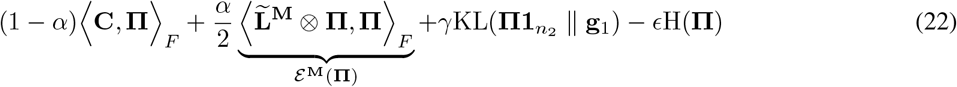

As indicated by the underbrace in (22), we define the *merged feature-spatial energy*, denoted *ε*^**M**^(**Π** ) to be the sum of 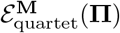 and 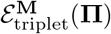, to have a more descriptive way of referring to the left-hand side of (21).

#### S1.3 Connection to energy-based models

Owing to the presence of multiple energetic conditions, including within-slice energies and matching energies, we make a connection between the optimal transport problem we have defined and an energy-based model. For some energy function *ε*_θ_(***x***) ≥0, and for some variable we are optimizing ***x***(≜ **Π** in our case), one may define a Gibbs-Boltzmann distribution *P*_θ_(***x***) given by:

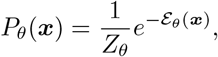

where *θ* comes from the SRT data, *θ* = (**C, C**^(1)^, **C**^(2)^, **X**^(1)^, **X**^(2)^). The normalizing constant (or *partition function*) *Z*_θ_ is given by 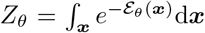, which is generally intractable to compute. For such an energy-based model, in

order to find an optimal value for ***x***, one may optimize the the log-likelihood of the model given by ∇_***x***_ log *P*_θ_(***x***), where:

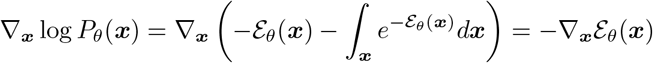

which holds as the partition function is clearly a constant with respect to ***x***. For the Wasserstein energy ⟨**C, *x***⟩ ,if *ε*_θ_(***x***) = −⟨**C, *x***⟩_F_ represents our energy function, we are simply solving a standard optimal-transport problem. If *ε*_θ_(***x***) = − ((1 − *α*) ⟨ ***C, x***⟩ _F_ + *α* ⟨ **L**^**D**^ ⊗ **Π, Π**⟩ _F_), then we are solving an FGW optimal-transport problem, and so on. While the connection to energy-functions is trivially evident, the value of introducing the connection is seen in the product-of-experts formulation of energy-based models, where one may have multiple Gibbs-Boltzmann distributions 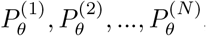, each carrying some “expert” information, where the distribution representing the combination of the features of all of the distributions is called a product-of-experts [16], given by:

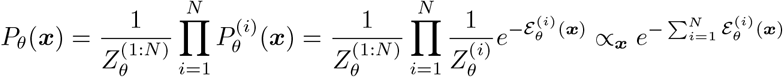

Where the energy-function of the final Gibbs-Boltzmann distribution is given by the *sum* of the individual energies:

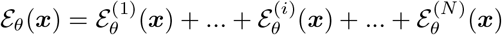

As such, we augment the optimal-transport formulation by framing it as a product-of-experts, constructing a total-energy which we optimize with respect to **Π** that carries the standard Wasserstein (doublet) energy ⟨**C, Π**⟩_F_, the quartet energy ⟨**L**^**M**^ ⊗ **Π, Π**⟩_F_ defined through the merged feature-spatial matrices, and two triplet energies which are also defined from these matrices: Tr **ΠM**^(2)^**Π** ^T^ and Tr **Π** ^T^ **M**^(1)^**Π** . Denoting the optimal-transport entropic-term density *P*_ℋ_, we minimize the following product-of-experts Gibbs-Boltzmann distribution with respect to **Π** :

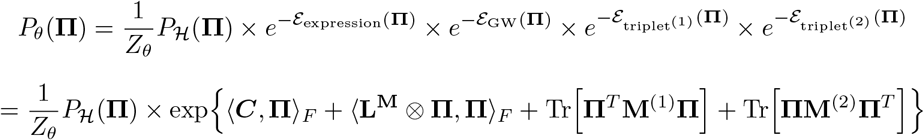

Which offers the interpretation that taking the sum of the energy terms in our optimal-transport problem is equivalent to optimizing a product of the Boltzmann-distributions over our variable **Π** , where this product represents a product-of-experts in which each “expert” distribution refines the matrix **Π** to be more biologically and spatially realistic with respect to the transcriptomic and spatial geometry of the first and second slices. Alternatively viewing each term as representing pairwise motifs (Wasserstein, or *doublet* term), ternary motifs (*triplet* term), and quaternary motifs (GW, or *quartet* term), these experts capture features ranging from lower to higher order.

### S1.4 On the mass-flux term, semi-relaxed OT

The semi-relaxed approach has been explicitly discussed in [41, 12, 6, 4, 32, 7]. Blondel et al. [4] formulate a version of our problem in their Definition 3 (“semi-relaxed smooth primal”), fixing their second marginal as we do, but without using entropic regularization in order to produce sparse transport plans. These authors point out that the mixed relaxed distance introduced by Benamou in [2] is a version of semi-relaxed OT, fixing the first marginal instead of a second, and using an *ℓ*^2^-penalty in place of a KL penalty. Interestingly, this penalty is applied directly to the analogue of ***ξ*** in this setting. A recent paper of Dong et al. [12] uses semi-relaxed transport for domain adaptation, aligning imaging data of different modalities. These authors use a Wasserstein loss, fixing the first marginal and in addition constraining the total mass transported. Vincent-Cuaz et al. [41] apply the semi-relaxed framework to graph matching, fixing the first marginal, and using a GW loss function. For matching two graphs, they find that fixing the first marginal while relaxing the second better preserves the structure of the first graph under the transport plan, versus the balanced regime. These authors [41] note that semi-relaxed OT (in the absence of entropy regularization) produces sparser transport plans than vanilla optimal transport. This sparsity also motivated the earlier work of [32] on color transfer; maps that are “too” multi-valued (i.e. not sparse enough) can inappropriately mix colors being transferred and create spatial irregularities. These authors also fix the first marginal, using a sparse transport plan to assign colors from the second image to the pixel of the first, while preserving its geometry. On the theoretical side, Chizat et al. [7] introduce the notion of a semi-coupling to connect the (PDE) dynamic and static pictures of semi-relaxed OT. Lastly, we mention two studies of Frank-Wolfe [13] and Sinkhorn [14] in the semi-relaxed setting.

We now discuss some properties of the mass-flux term ***ξ*** built from **Π** and advantages our formulation give us. Let us start by examining the objects analogous to ***ξ*** in the fully-unbalanced case. When both marginal constraints are relaxed, both of the vectors 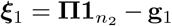 and 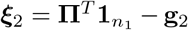 may be non-zero. Towards relating vectors ***ξ***_1_ and ***ξ***_2_, we take the inner product of these expressions with 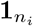 to obtain:

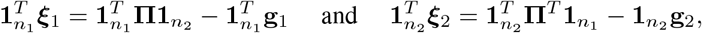

We solve each of these for the total mass transported, namely 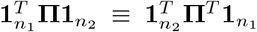. After relating the two expressions through the total mass transported and re-arranging terms, one has

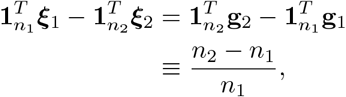

with the last line following from our choice of normalization on the uniform measures 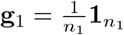 and 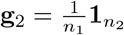. In the semi-relaxed context of `DeST-OT`, ***ξ*** ≡ ***ξ***_1_, and vector ***ξ***_2_ is zero, in which case, one has

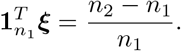

Thus, the semi-relaxed formulation allows us to describe all growth and death implicit in transport plan **Π** by a single vector ***ξ***, instead of a pair of vectors (***ξ***_1_, ***ξ***_2_) of different dimensionality. Moreover, the hard constraint of the semi-relaxed formulation makes this single vector ***ξ*** interpretable in a way that the pair (***ξ***_1_, ***ξ***_2_) is not. Observe that the second marginal **g**_2_ is fixed, with units assigning mass 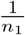 to each spot. Though the first marginal of **Π** is no longer fixed, ***ξ*** is by definition an additive perturbation of **g**_1_ which uses the same units as **g**_2_, also assigning mass 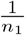 to each spot. ***ξ*** also uses these units, giving expression **g**_1_ + ***ξ*** physical meaning.

Rescaled vector 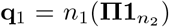 has in its *i*-th element the amount of mass transported from spot *i*, in units of *numbers of spots*. These numbers are fractional, as **Π** is implicitly estimating these values in a sense averaged over *t*_2_ −*t*_1_, with respect to a stochastic process similar to the one described in [6]. Because the rescaled target measure *n*_1_**g**_2_ shares these counting-measure units, we can reasonably interpret **q**_1_ ≡ **q**_1_(**Π** ) as the vector of the *expected number of descendants* of each spot in the first slice, as predicted by **Π** . This is clear, as we have:

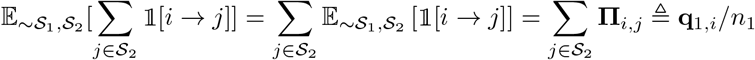

Where multiplying by *n*_1_ scales **q**_1_ to have the units of counting measure over spots. We lose this interpretation in the fully-unbalanced regime: when the target measure is not constrained to be uniform, the amount of mass transported *from* spot *i* may not reflect an expected number of descendants, as this value will depend on the mass of each descendant.

From the definitions of ***ξ*** and **q**_1_, we have

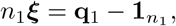

and *n*_1_***ξ*** has the following intuitive interpretation: at spot *i*,

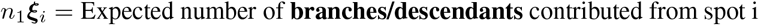

## S2 Growth rate conversion

The conversion of the growth vector (mass-flux) ***ξ*** to a growth rate can be done simply. We use the growth process described in [22], where one considers the PDE modeling a transcriptomic trajectory as a density evolving in time:

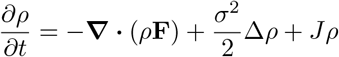

With −**∇ ·**(*ρ***F**) a continuity equation drift term, 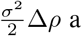 a diffusion term, and *Jρ* a term representing growth and branching, which are all modeled using successive optimal transport steps. We convert our mass-flux term from the optimal transport to a growth using the principle of [22], where operator-splitting the branching component above from the drift-diffusion gives us a continuous-time growth update as:

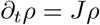

Thus we simply need to consider the solution to the ordinary differential equation 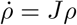 for *J* between times *t*_1_, *t*_2_:

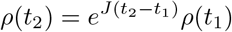

For *ρ*(*t*_1_) and *ρ*(*t*_2_) a prior and posterior density over the spots in the first slice. By the semi-relaxed optimal transport formulation, we simply compare against a simple uniform prior density 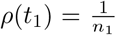 which we start with. Thus one concludes the connection to a growth rate, on a per-spot basis in the first slice, is given as:

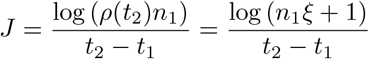

Which establishes a simple monotonic connection between our growth vector ***ξ*** and a proper growth rate in time. In the notation of [22], *τ* = *t*_2_ −*t*_1_, and one has *J* = *τ* ^−1^(*p*_b_ −*p*_d_), establishing a connection between the growth rate and birth-death parameters *p*_b_, *p*_d_, and *τ*. Conventionally, cell division and death are modeled as follows: each cell has an exponential clock of rate *τ* ^−1^. When a cell’s clock rings, it dies with probability *p*_d_, or splits into two cells with probability *p*_b_ = 1 −*p*_d_. As such, our growth rates have a natural connection to the parameters of this birth-death process.

## S3 Connection to the continuity equation and interpretation of growth vector

Suppose we denote **x**(*t*_k_) = **x** ∼*μ*, **x**(*t*_k+1_) = **y** ∼*ν* as samples of our slices 𝒮_1_ and 𝒮_2_, and consider the pairwise-alignment problem in the equivalent Monge-formulation (the discrete analogue of *μ* being **g**_**1**_, and *ν* being **g**_**2**_). The Monge-problem is formulated as:

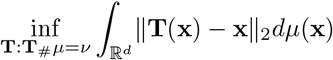

Where it can be shown that an equivalent form exists in which the transport-map can be expressed as the integral of some vector field **F**(**x**, *t*), weighted by a time-dependent density *ρ*(**x**, *t*) and where the optimization is performed over a vector-field of least-norm:

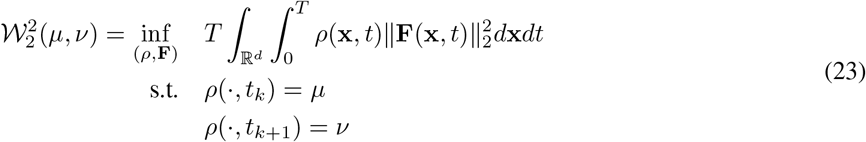

Where the vector field is optimized subject to the continuity equation: *∂*_t_*ρ* = −**∇ ·**(*ρ***F**(**x**, *t*)) [39]. In the semi-relaxed case, one may either assume mass-conservation or generalize the vanilla continuity equation by assuming the presence of a source or sink term *σ* (i.e. *σ* = *Jρ* for *J* a growth rate as defined in S2) which allows mass to be introduced into the system in the form *∂*_t_*ρ* = −**∇ ·**(*ρ***F**(**x**, *t*)) + *Jρ*. As such, ***ξ*** represents the integral of a divergence in the case of mass-conservation (i.e. local mass-redistribution), or a spatial divergence added to a source/sink term in the case where the total mass is increased or decreased in the system. To be precise, we have that in the balanced case:

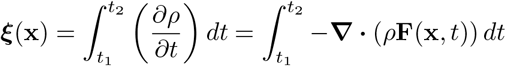

And general semi-relaxed case:

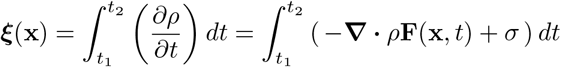

## S3.1 Interpretation of *ξ*

Supposing we call the spatial volume encompassed by our tissue slice at time *t, V*_t_. Each element of *ξ, ξ*_k_, represents the approximate value of **∇ ·**(*ρ*(*x*_k_, *y*_k_)**F**(**x**, *t*)(*x*_k_, *y*_k_)) in some box [*x*_k_ − Δ, *x*_k_ + Δ] × [*y*_k_ − Δ, *y*_k_ + Δ] centered at (*x*_k_, *y*_k_) ∈ *V*_t_. Generally speaking the volume *V*_t_ is assumed to be connected such that, for *N*_t_ dots at time *t*, the volume is 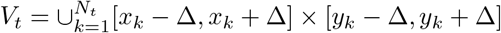.

We assume a cell type assignment is some function from a matrix of distance-based dissimilarities and feature-values to a partition 𝒫 of the data *ϕ*(*D, C*) = 𝒫. Where 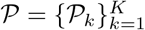 represents a decomposition of our dots into disjoint sets representing respective cell types. Following from this, we define the volume of a cell type *k* at time *t, V*_t,k_, as 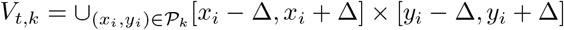. Clearly, 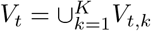.

### Definition 1.

*Tissue-slice growth. The term* 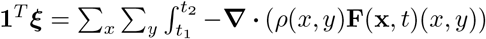 *represents the discretized volume integral*

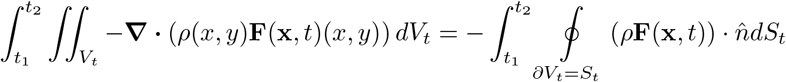

*for* Δ = 1*/*2. *By the divergence theorem, one identifies the sum of the divergences in the volume V*_t_ *as the mass-flux across the boundary ∂V*_t_. *Thus, the growth of a tissue slice contained within in a volume at time t, V*_t_, *is defined as the mass-flux across this boundary integrated from t*_1_ *to t*_2_.

For a balanced optimal transport this is identically zero for any boundary as ***ξ*** = **0**:

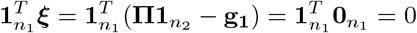

For a semi-relaxed optimal transport with marginals that sum to the same value, such as the uniform marginal 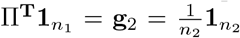 and marginal 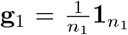, one may have *local* growth or death as ***ξ*** is not necessarily the zero-vector, but globally mass is conserved:

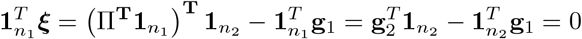

In the case of a semi-relaxed optimal transport, if we allow for this source or sink term where

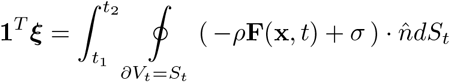

we generalize the above definition by asserting global mass gain or loss is possible. In particular, we can set the unbalanced marginals to reflect the normalized change in volume (or change in mass) as:

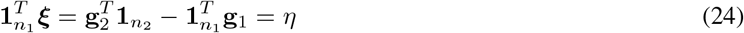

Where *η* ∈ ℝcould be chosen to reflect something known–such as a total change of mass or volume between the two slices. We also note that the interpretation of ***ξ*** can also be used to measure local growth of a tissue within some fixed boundary, as described in the corollary below.

### Corollary 1

(Cell type growth). *Given a partition* 𝒫 *of tissue into separate cell types, and defining the vector* **z**_p_ *by*

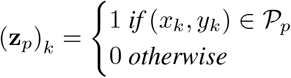

*The term* 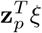*represents the volume integral*

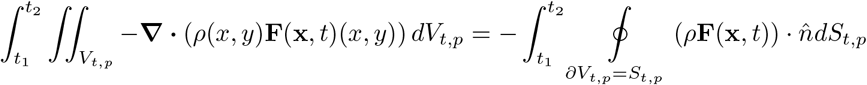

*and represents the mass-flux out of the boundary of cell type p, ∂V*_t,p_, *integrated from t*_1_ *to t*_2_.

### S3.2 Defining an Unbalanced Optimal Transport problem to match an a priori known total mass-flux

We seek a balancing condition which fixes the mass-flux *η* from our optimal transport 24 to some interpretable and meaningful value. By our previous decomposition of the volume of a bar-coded grid of spots into boxes of equal volume Δ^2^, or analogously a small hexagonal volume for Visium, and our assumption that the mass-density is constant for each spot, we have that the normalized mass-flux above must equate to:

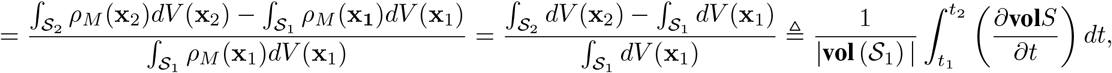

as we assume that the mass-density *ρ*_M_ (**x**) = *C* is constant for any spot on the grid.

For Δ^2^ a per-spot volume, we see that the volume-normalization lets this technology-specific factor cancel and the sum of the growth ***ξ*** is related to the change-in-spots directly as

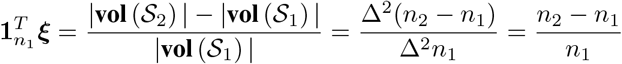

Which requires that the unbalanced condition have 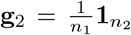 as the constraint measure. This makes sense, as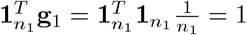 and 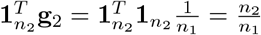, which has the interpretation that the density in the first slice sums to 1, and the density of the second is proportional to the ratio of the number of spots of the second slice to the first. With 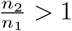 implying mass is generated as the second volume exceeds the first, 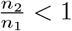 implying mass is lost as the second volume is smaller than the first, and 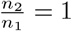 implying mass-conservation.

## S4 Semi-relaxed Sinkhorn

### Calculating the gradient with respect to Π

In order to run Sinkhorn, we require an initial condition on **Π** and an analytical expression for the gradient of our primal objective with respect to **Π** . In particular, the following represents a primal feasible **Π** :

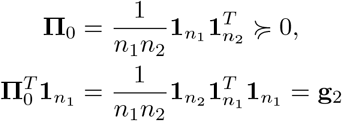

Recall that the energy function ε : **Π** 1→ ε (**Π** ) of interest in our current formulation is

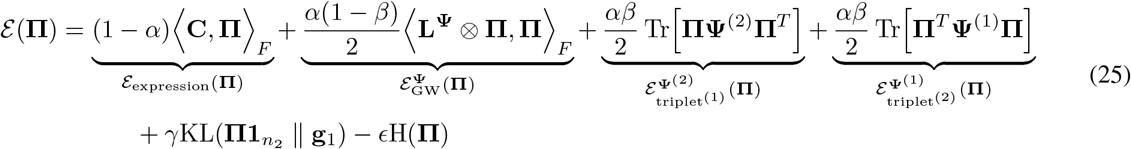

We calculate the gradient of *f* as follows. The gradient of the expression term ε_expression_(**Π** ) is straightforward to compute:

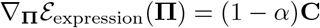

Next, we consider the gradient of the distance-regularization term E_row_ corresponding to the rows of **Π** :

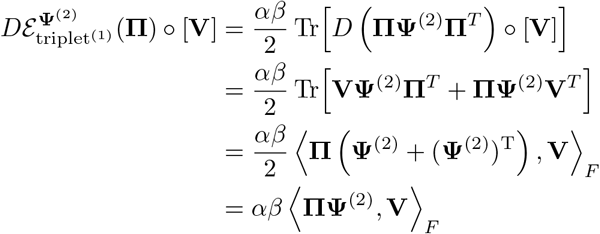

So, we have that:

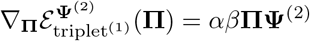

The gradient of the distance regularization term corresponding to the columns of **Π** is similarly given as:

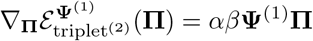

We now compute the gradient of *f*_KL_. Let 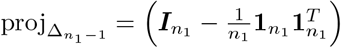 denote the projection onto the probability-simplex 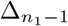

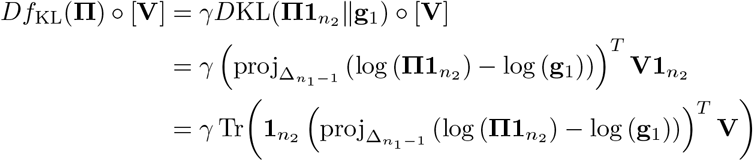

With, and the log applied element-wise above. So:

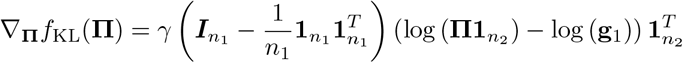

Lastly, we compute the semi-relaxed gradient for *f*_FGW_(**Π** ):

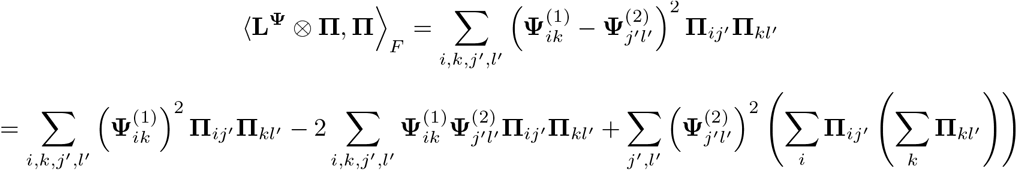

Owing to our semi-relaxed marginalization constraints, it holds that 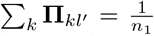 and 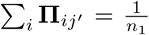, such that the rightmost term is a constant and disappears when we take the gradient with respect to **Π** . As such, the above is proportional to:

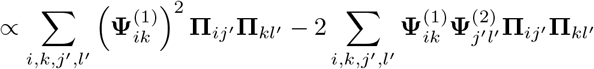

Converting this to matrix form, we have:

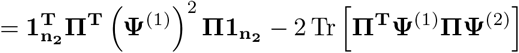

We consider each term independently. For the rightmost we consider the Fréchet derivative in the direction **V**:

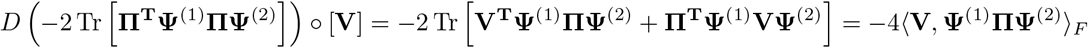

For the leftmost term, we have:

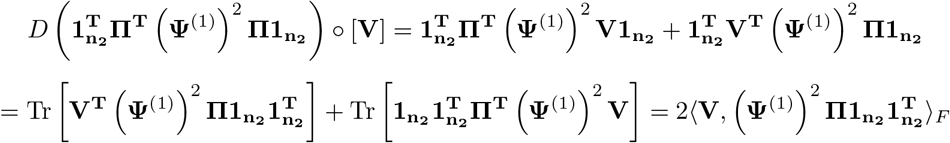

We can therefore identify the final gradient of our semi-relaxed GW term as:

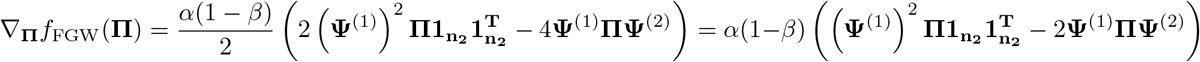

With the squaring operation applied element-wise through **Ψ**^(1)^.

### S4.1 Dual formulation of semi-relaxed Sinkhorn

For our optimal-transport problem, we have both a primal objective E(**Π** ) we seek to minimize, as well as a single constraint that 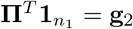 which restricts the set of primal-feasible **Π** . The Karush-Kuhn-Tucker (KKT) conditions introduce a set of dual variables for each constraint, and a Lagrangian which lower bounds the primal objective. The conditions, which involve first order constraints, dual constraints, and complementary slackness constraints, offer a set of conditions which must be satisfied by an optimal solution to the primal problem. As such, we introduce the dual-variable ***μ***, and establish the Lagrangian ℒ to maximize as a lower-bound to the primal objective:

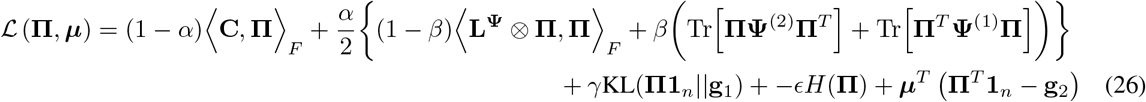

Collecting the non-entropic terms in the gradient of the above expression, we define the matrix **C**^*^ to be:

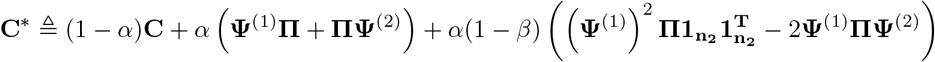

The KKT conditions imply the following first-order condition holds when we consider the gradient of our Lagrangian with respect to the variable being optimized, **Π** :

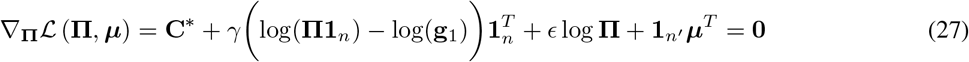

We also have the following primal constraint for our singular equality condition on the second marginal:

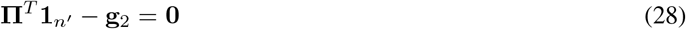

The dual constraints, which are implicitly accounted for in the entropy term of Sinkhorn which lifts the matrix **Π** through exponentiation such that **Π** ≽ **0** always holds, for technical correctness include a dual variable **Ω** ≽ **0** and a term in the Lagrangian of the form −⟨Ω, **Π** ⟩_F_. The two dual constraints would formally be given as:

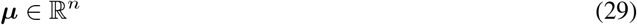

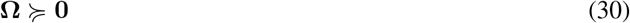

And a complementary slackness condition enforced elementwise through the matrix **Π** , where one either has the active constraint where **Π** _ij_ *>* 0 and **Ω**_ij_ = 0, or a slack, or inactive, constraint with **Ω**_ij_ *>* 0 for the (omitted) component of the Lagrangian given as −⟨**Ω, Π**⟩_F_. Thus the slackness condition would be written as:

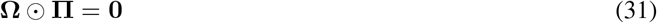

Looking more closely at the first-order conditions, one finds:

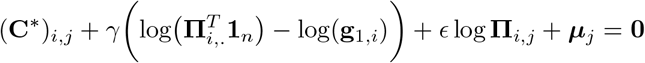

Implying a similar solution-form to Sinkhorn involving an exponentiation with the Gibbs kernel, wherein one has 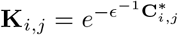 and:

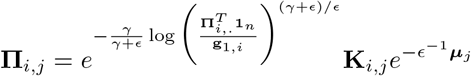

From Sinkhorn, we know there exist unique vectors **u, v** such that **Π** = diag (**u**)**K** diag (**v**) and see:

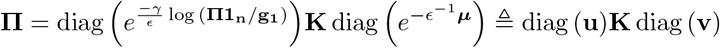

We consider what the update rule should be, given this identification, by establishing:

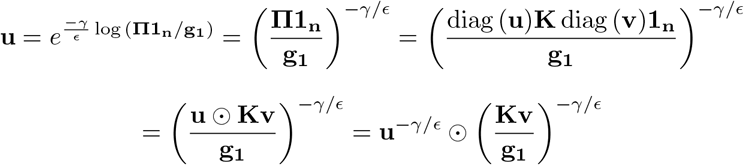

Noting the Hadamard multiplication, this directly implies an update rule for **u** in the form:

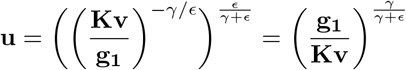

This may analogously be identified from the general form of unbalanced optimal transport, wherein one uses an anisotropic proximal step weighting two divergences on the marginals **g**_**1**_. In particular, for a generalized optimal transport problem of the form:

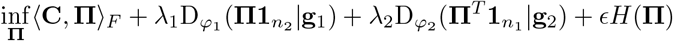

for *φ*_1_, *φ*_2_ convex, positive lower-semi-continuous divergences. For these, one takes a proximal step for each marginal where **u, v** are updated as:

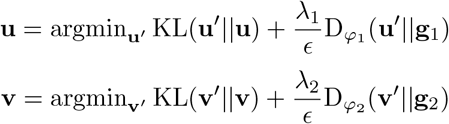

This has the following closed-form updates for unbalanced Sinkhorn where we consider the case that 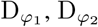 are taken to be KL-divergences:

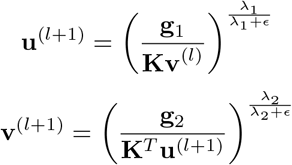

While we demonstrated that the semi-relaxed update follows directly from the KKT conditions, one can easily recover the semi-relaxed update in the limit of *λ*_2_ → ∞ in the unbalanced case. The marginal update then becomes 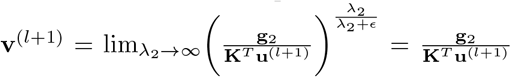, where one recovers the marginal constraint on **g**_2_ and our semi-relaxed update rule.

### S4.2 Converting semi-relaxed Sinkhorn to log-domain for stability

Consider a general form of entropy-regularized, fully unbalanced Wasserstein optimal transport: given measures 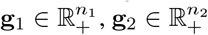, and given a candidate transport plan 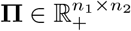, let the functional **Π** 1→ P_ϵ_(**Π** ) be given by

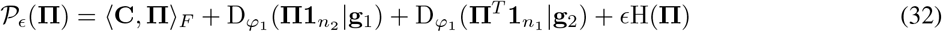

where *φ*_1_, *φ*_2_ are convex, positive lower-semi-continuous, and with *φ*_i_(1) = 0. The terms 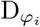 are the associated *φ*-divergences. Depending on the choice of the *φ*_i_, one can recover either the fully unbalanced constraints of moscot, or the semi-relaxed setting of `DeST-OT`. Specifically, `moscot` corresponds to taking *φ*_1_ = *φ*_2_ ≡ *p*→ *p* log *p* −1,*p* + 1, in which case both *φ*-divergences are the KL divergence. On the other hand, `DeST-OT` corresponds to taking *φ*_1_ ≡ *p* 1→ *p* log *p* − *p* + 1, while the single hard constraint corresponds to choosing *φ*_2_ = *ι*_{1}_, where

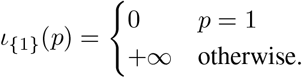

Thus, (32) corresponds to a version of the `moscot` objective (3) with the GW term ⟨**L** ⊗ **Π, Π**⟩ omitted, or a version of the `DeST-OT` objective (22) without our modified GW term 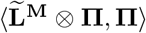. In both cases, the hyperparameters attached to different terms have been suppressed.

We define 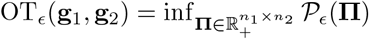, and we appeal to the dual formulation of (32), which we state as a proposition.

#### Proposition 1

([36], Proposition 2). *One has*

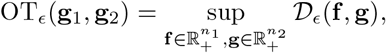

*where for such* 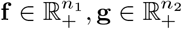, *we define the dual objective to* 𝒫_ϵ_ (32)

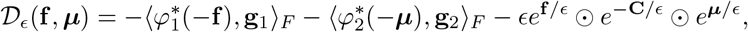

*where* 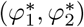 *are the Legendre transforms of* (*φ*_1_, *φ*_2_), *namely* 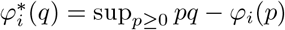

The optimal primal plan **Π** ^⋆^ is obtained from the optimal pair (**f** ^⋆^, ***μ***^⋆^) of dual variables as

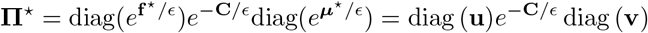

Thus, connecting the dual variables of the generalized Sinkhorn objective for arbitrary *φ*-divergences to our primal variables **u, v**, we are able to express closed-form updates for the dual variables instead. The Sinkhorn iterations for the semi-relaxed case, in terms of the dual variables **f**, **g** are given as:

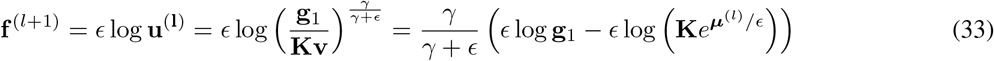

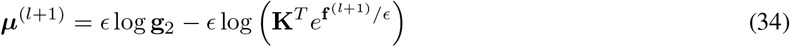

Owing to the presence of the exponential of each dual variable on the right-hand side of 33 and 34, these cannot be evaluated directly for small values of the entropy-regularization *ϵ*. When *ϵ* →0 we would recover an exact, non entropically-regularized optimal transport – however, this would cause the term inside of the logarithm to exceed numerical precision. For the purpose of introducing the log-domain Sinkhorn algorithm, we make the following definitions. For 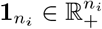 and *ϵ >* 0, the *Softmin operators* 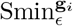 are defined for any vectors 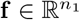 and 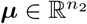 by:

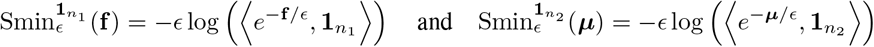

Rewriting 34 34 we see:

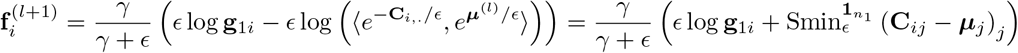

And:

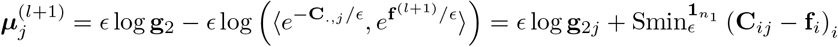

One can simply evaluate the softmin 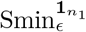 using the log-sum-exp trick, which we broadcast across dimensions for an efficient and stable log-domain update.

## S5 Metrics of growth

Let 𝒮_1_ = (**S**_1_, **X**_1_) and S_2_ = (**S**_2_, **X**_2_) be a pair of spatiotemporal tissue slices, corresponding to timepoints *t*_1_ and *t*_2_. For a collection of cell types shared across both slices, enumerated as {1, …, *P* }, suppose we are given cell type partitions 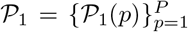 and 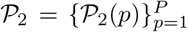for each slice. The *mass of cell type pat time t*_1_ is *m*_1_(*p*) = |𝒫_1_(*p*) |, and the *mass of cell type pat time t*_2_ is likewise *m*_2_(*p*) = |𝒫_t_(*p*) |. The *mass-flux of cell type p* over these two timepoints is then given by *m*_2_(*p*) −*m*_1_(*p*). Below, we define a metric quantifying how well the cell type mass-flux inferred by an optimal transport method matches the empirical mass-flux of a cell type. If such a method outputs a pair (**Π, *ξ***) of an alignment matrix and a growth vector, we measure the distortion between ***ξ*** and the true mass flux. We discuss the assumptions that go into this metric, and how it can be generalized.

### S5.1 A metric of growth: individual cell level

To measure the accuracy of the growth rates across cell types *and* spots within the cell type, we define the true growth rate at time *t*_1_ of cell type *p* as:

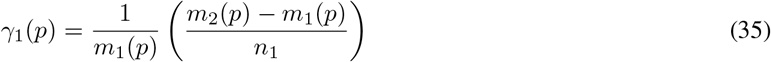

In calling this a true growth rate, we are making two assumptions: first, that there are no cell type transitions between distinct cell types – in the next subsection we discuss how to relax this. The second assumption made is that the burden of accomplishing the change in mass is shared equally across cells of the same type, which can be viewed as an entropy-maximizing assumption.

The normalization factor 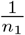 makes these growth-rates consistent with the slice one mass-normalized values of the marginals **g**_**1**_, **g**_**2**_ and ensures consistency across experiments. We quantify the total distortion between the true growth factors for each cell type and those derived from optimal transport as:

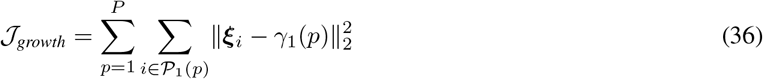

This assesses the total divergence between the optimal-transport growth rates and the true, empirical growth rates under a fixed cell type labeling. The sum of the growth-rates across spots within a tissue slice equals the total growth rate of the slice:

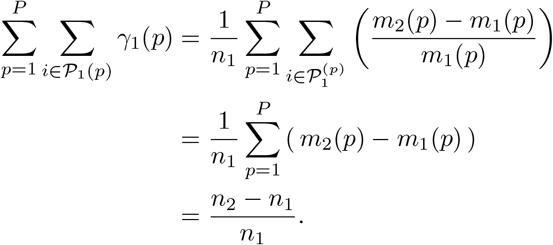

### S5.2 Generalizing the growth-metric to cell type transitions

The definition of growth given in the section above assumes that cells of type *p* transition and grow to their own cell type, and computes the distortion relative to this assumption. While this is valuable as a baseline metric when no cell type transitions are available, we generalize this notion when we have a density matrix of cell-to-cell reverse-time transitions 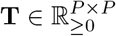 across *P* cell types. Supposing we have a vector 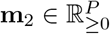of cell type masses at time *t*_2_, and likewise **m** _1_ at time *t*_*1*_, we modify *m*_*1*_ (*p*) in the definition above to include transitions. The section above implicitly assumes that the matrix is diagonal (in particular, it is the *P* ×*P* identity matrix 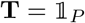). Under ground truth transition matrix **T**, the mass of cell type *p* from time at time *t*_1_ is described in terms of **m**_2_ as follows:

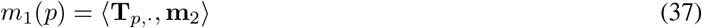

Which is to say, **m**_1_ can be described in terms of **m**_2_ via the matrix-multiplication

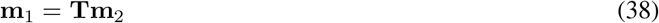

and the mass-flux out of cell type *p* remains as defined in the previous sections with the only adjustment being that the new *m*_1_(*p*)’s account for cell type transitions. As before, we assume the cell type partitions across each slice are consistent with these cell type mass vectors.

Now, consider an alignment matrix 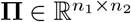, and let {*p* → *q*} denote the following set:

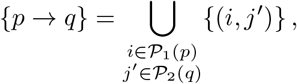

understood as the event that cell type *p* of the first slice is mapped to cell type *q* in the second slice. In making this definition, we show how any alignment matrix **Π** induces a reverse-time transition matrix: indeed, **Π** is a positive measure on such pairs, so one can naturally form a coarse-grained *P* ×*P* matrix from **Π** using the events {*p* →*q}* : first, define

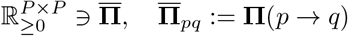

Whatever **T** we construct from **Π** should be a transition matrix, row-normalized such that **1**_P_ = **T1**_P_. This is achieved by tilting each row *p* of measure 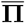 with the factor *n*_1_*/m*_2_(*q*):

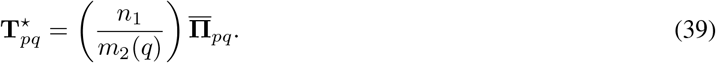

Now, consider the distortion metric defined in (36). Instead of viewing 𝒥_*growth*_ as a function of ***ξ*** only, as was done in the previous section, we now regard it as a function of both ***ξ*** and 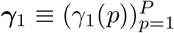. Just as ***ξ*** arises from **Π** , vector ***γ***_1_ is defined from **T** through (37) and (35). Given the output (**Π, *ξ***) of an OT method, the next proposition gives the optimal pair (**T, *γ***_1_) for this output according to the growth distortion metric (36).

#### Proposition 2.

*Given cell type partitions 𝒫*_1_ *and 𝒫*_2_ *of two SRT slices, let* **m**_1_ *and* **m**_2_ *be a pair of cell type mass vectors consistent with these partitions, and related by a transition matrix* **T** *through* (38). *Fix an alignment matrix* **Π** *with* 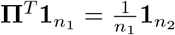 *and consider its associated growth vector* ***ξ*** ≡ ***ξ***(**Π** ). *Then, for* ***γ***_1_ ≡ ***γ***_1_(**T**), *one has*

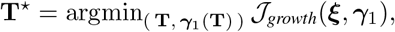

*with* **T**^⋆^ *defined from* **Π** *in* (39).

*Proof*. When ***ξ*** is fixed, the minimizer of 36, in terms of ***γ***_1_, is given in each entry by the cell type sample mean of ***ξ***:

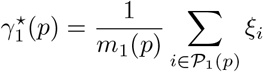

Thus, we evaluate the right-hand side to find:

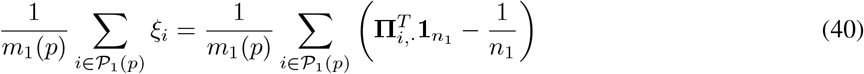

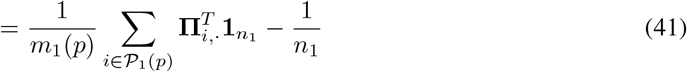

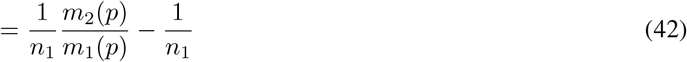

Therefore, we have that:

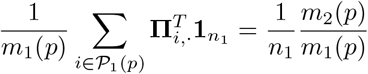

And we decompose the following sum according to the cell types *p* may transition to in the next slice:

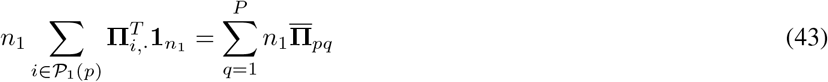

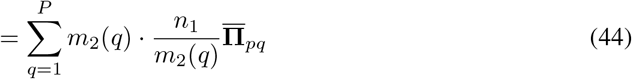

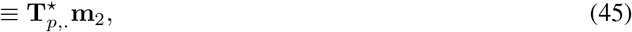

which shows that the optimal 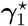 arises from the claimed optimal transition matrix **T**^⋆^ defined in (39) We complete the proof by checking that each row of **T** sums to one. For notational convenience let **z**_2_(*q*) denote the vector in 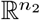whose *j*^′^-th component has a 1 if and only if *j*^′^ ∈ 𝒫 _2_(*q*), and is 0 otherwise; likewise, let **z**_1_(*p*) be the vector in 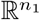 which is a one-hot encoding of cell type *p* over the first slice.

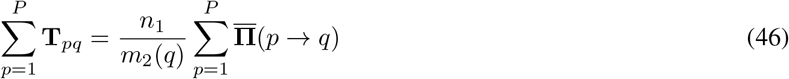

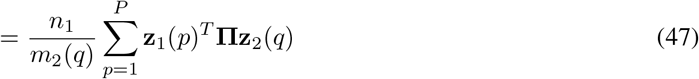

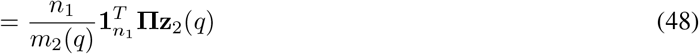

The last step is to use the single hard constraint in our semi-relaxed formulation, along with our choice of mass calibration for the second slice: 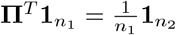. Continuing from the above display, one has

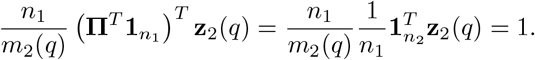

By definition, all entries of **T** are necessarily positive. Therefore these row-sum conditions show us this is indeed a reverse-time transition matrix, and the proof is finished.□

### S5.3 Interpretation of per-cell growth distortion 𝒥_growth_

For the setting of *γ*_1_(*p*) in the previous section, rather than matching the cell type mass-fluxes assuming cell types transitioning to themselves, we find the optimal matrix **T** of cell types transitioning from *t*_1_ to *t*_2_, calculate the adjusted mass contributed to **m**_2_ from cell type *p* at *t*_1_ as *m*_1_(*p*) = **T**_p,·_**m**_2_, and calculate the expected distortion in the mass-flux. As noted in 2, the minimizer of 𝒥_*growth*_ over ***γ***_1_ is simply the sample mean of the per-cell type mass-flux, given as:

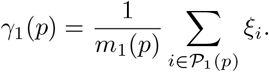

Therefore an alternative interpretation to one in which we match the mass-flux of the optimal transition matrix is that we are measuring the sum of the variances of each mass-flux variable within each cell type:

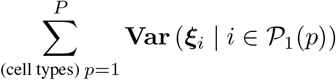

In a tissue or cell type, the assumption that growth is relatively homogeneous or smoothly varying, as opposed to coming from a very small, sparse, set of cells, has been observed in morphogen studies where within tissues like the imaginal wing disk in drosophila one observes uniform cell divisions and growth [15] [25] [27]. As such, measuring the within-cell type variance of the growth-factors is biologically reasonable and captures whether growth patterns are very sparse (i.e. when aligned via an unbalanced routine which assigns a few “representative” cells as those which grow), or whether growth patterns are relatively consistent.

## S6 Procrustes-like problems and a cell migration metric

### S6.1 Quantifying a metric of naive rigid-body migration

Suppose we are given the solution to either the generalized Procrustes’ problem for an orthogonal transformation **Q** ∈ O(*n*) or the generalized Wahba’s problem for a rotation **Q** ∈ SO(*n*) along with a general translation vector **h** ∈ ℝ^n^ as described in S6.3. This then describes a simple rigid-body transformation which relates the coordinate frames of slice 1 and 2, 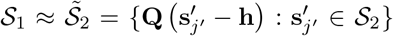. Supposing the true transformation relating the two frames is indeed a rigid-body transformation, one may directly quantify the cost of an alignment with respect to the matrix **Π** given **Q, h**. We introduce this as our *naive rigid-body migration* metric, given as:

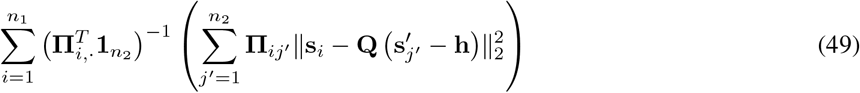

Where we take the difference between the spatial points **s**_i_ and 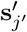 in our pair of slices (**X**^(1)^, **S**^(1)^) and (**X**^(2)^, **S**^(2)^). This difference is computed under the posterior implied by **Π** , and as we compare matrices **Π** which may be balanced, unbalanced, or semi-relaxed, we normalize by a factor of 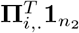 so that any posterior in 49 is normalized to a distribution consistently. We call this metric *naive*, as it is clear this transformation has determinant 1, and neither allows for volume expansion (growth) or volume shrinkage (death). Moreover, the true transformation underlying tissue development might involve some arbitrary (e.g. diffeomorphic) transformation, such that the rigid-body assumption is more generally inappropriate. As such, it is not necessarily the case that lower loss under this metric is universally better. However, one can say that if the alignment tends to incur exceptionally high loss under this metric, without any geometric consistency, the alignment may be spatially unrealistic. In other words, it might be aligning the sub-level set of points with similar features 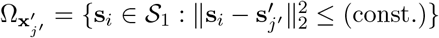 without accounting for the geometry of either slice.

### S6.2 Invariance of the method to rigid-body transformations, or group action by members of SE(2)

Let us call each spatial point in the first slice **s**_i_ ∈ ℝ^2^, and each point in the second slice 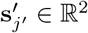. Suppose we want the second slice to be in the same coordinate basis as the first slice, but we do not know **Q** ∈ SE(2), **h** ∈ ℝ^2^, representing the rigid-body transformation of the data 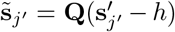 which moves us into the correct, shared coordinate-frame. A reasonable question to ask is whether our objective returns an alignment matrix **Π** which is unique, independent of the specific **Q, h** that relate the two coordinate frames. From the general objective in 20, it is clear that our objective only depends on the points in the first and second slices 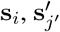 through the distance matrices **D**^(1)^ and **D**^(2)^. Considering that **s**_i_ is in the correct basis, let us restrict our attention to 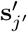 and **D**^(2)^. It is simple to check that:

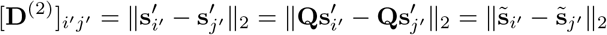

Which implies that our objective does not depend on the spatial location or rotation of the second slice relative to the first. Uniqueness therefore only depends on the convexity of the objective, which could in principle be achieved by using the projection of **D**^(1)^, **D**^(2)^ onto the vector space of positive-semi definite matrices 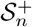 or kernelization.

The next reasonable question is whether one can recover the correct **Q, h** given the alignment matrix **Π** . Problems of this form include the Orthogonal Procrustes problem (where **Q** is simply an orthogonal matrix), and Wahba’s problem (where **Q** ∈ SO(2) explicitly), which are both well studied and have existing solution methods. Thus, the final problem needed to align the coordinate frame can be given (in Procrustes’ form) as:

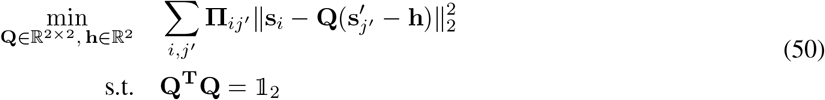

Or, in Wahba’s problem form:

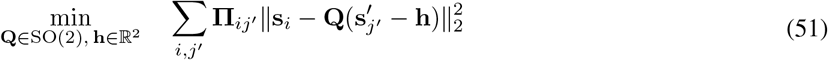

We offer both the Procrustes’ and Wahba’s form, with the latter having the restriction that **Q** ∈ SO(2) requires det **Q** = 1, which is to say we seek an explicit (orientation-preserving) rotation rather than a general orthogonal transformation (which may be a rotation composed with a reflection). In either case, the problem of finding a correct alignment is something that emerges *after* the outputs of `DeST-OT`, which is itself completely independent of the coordinate frame the first and second slice are set in and is invariant to linear translation and rotation, unlike works involving Gaussian processes such as [18].

### S6.3 Review of the Solution to Wahba’s problem

First, we consider an appropriate transformation of the coordinates to eliminate the dependence on the translation, for this light generalization of Wahba’s problem to a joint-distribution across both slices [10]. In particular, consider the change of variables given by:

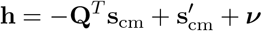

For 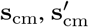 representing the center of mass of slice 1 and slice 2, and ***ν*** an undetermined constant. In particular,

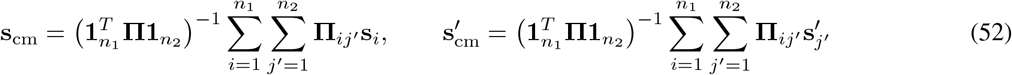

Substituting this into the primal objective:

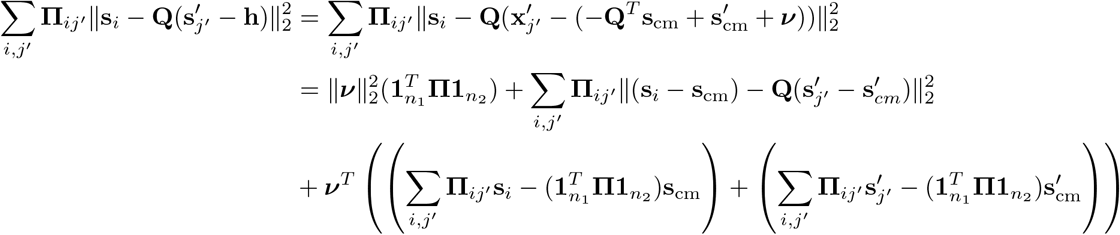

By definition of the center of mass-coordinates, the above equates to the following, with the minimization over **r** now over ***ν***:

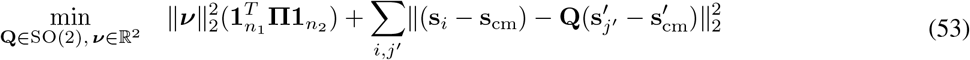

Where the independence of the left and right primal objectives yields that the minimizer is simply ***ν***^⋆^ = **0**. After the optimization for the rotation, the optimal translation is simply: 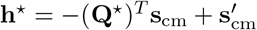. Thus, one may simply focus on the problem:

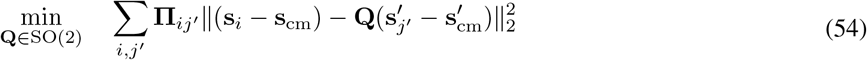

The presence of the joint distribution keeps this from being set in the standard form which Wahba’s problem is presented and solved in. We give a simple equivalent form, introducing the center-shifted coordinate matrices across spatial locations in the slices:

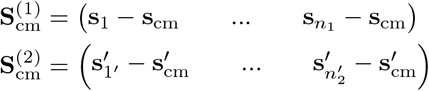

And consider the equivalent objective, with vec denoting the vectorization operation which column-stacks a matrix, **Π** ^1/2^ denoting the element-wise square-root of the entries of **Π** , and ⊙ denoting the elementwise, or Hadamard, product:

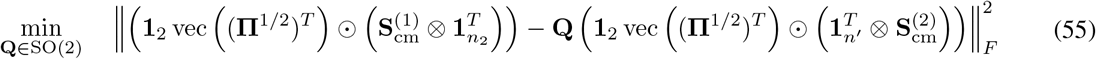

We rename the matrices to be concise as **G**^(1)^, **G**^(2)^:

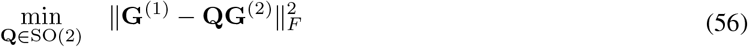

The simple solution [10] can be given using an SVD of **G**^(1)^(**G**^(2)^)^T^ = **UΣV**^T^. Where the final orthogonal transformation would be given as **Q** = **UV**^T^. If one adds the restriction that det **UV**^T^ = 1 for a rotation, one would find **Q** as:

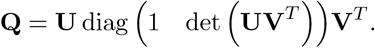

### S7 Generate feature (expression) vectors for simulated data

## S7.1 Generate feature vectors for simulated 1D slices

Our simplest experiment considers a pair of *one-dimensional* tissue slices with the same number of spots (51 total). Writing *N* in place of 25 for clarity, the spatial coordinates of the first and second slices are given by identical matrices, namely **S**^(1)^ ≡ **S**^(2)^ = [−*N* (*N*− 1) … −1 0 1 … (*N* −1) *N ]*, understood as a column vector. Synthetic features are constructed by first assigning one of two cell type labels (“A” or “B”) to each spot, and then assigning numerical features based on cell type. As in 2.3.1, cell types partition each domain, we write these as 𝒫_1_ and 𝒫_2_, where:

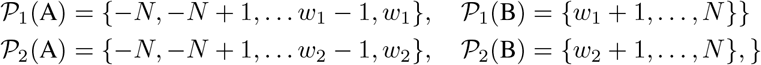

setting *w*_1_ = −10 and *w*_2_ = 5 in the experiment to be the “pivots” marking a change in cell type.

We generate eight-dimensional feature by sampling random vectors ***v***_1_, ***v***_2_, ***v***_3_, ***v***_4_ independently and uniformly from the unit sphere of ℝ^4^. Let **0 ℝ**^4^ be the zero vector, and let denote concatenation of vectors. We set 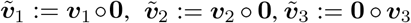 and 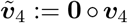. Over cell type A, we generate features by linearly interpolating feature 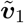 at spot *N* with feature 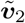 at spot *w*_1_ or *w*_2_, depending on the slice. Over cell type B, we generate features by linearly interpolating feature 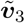 at either *w*_1_ or *w*_2_, with feature 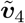 at spot *N*. Thus, each cell type is characterized by a feature gradient in four dimensions, and the features at distinct cell types are orthogonal by design.

### S7.2 Generate feature vectors for simulated 2D slices

We generalize the above by defining a two-dimensional model of tissue slices. Let *E* denote the centered, circular ellipse of radius *r* = 25:

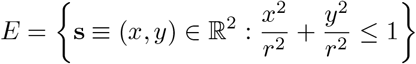

Let 𝕋 be the subset of the integer square lattice ℤ^2^ consisting of all integer pairs whose sum is odd: 𝕋= {(*x, y*) ∈ ℤ^2^ : (*x* + *y*) mod 2 = 1}. Then 𝕋 is a triangular lattice, mimicking the spatial organization of SRT data output by the Visium platform. Let **S**^(1)^ = **S**^(2)^ = **S** denote the stack of spatial coordinates associated to the set *E* ∩ 𝕋.

As above, we use a simple spatial condition to partition each slice into two pieces. Here, we set two “pivot” lines, given by *y* = *w*_1_ and *y* = *w*_2_. The cell type partitions 𝒫_1_, 𝒫_2_ over the two slices are defined as follows:

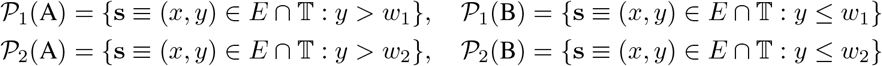

for *w*_1_, *w*_2_ = ±10.

We assign features in a similar manner to the one-dimensional case. Previously we assigned features to the boundary of cell type segments and linearly interpolated between these.

In this slightly more general setting, let *R*_*i*_(A) and *R*_*i*_(B) denote (smallest) bounding rectangles for sets 𝒫_*i*_(A) and 𝒫_*i*_(B) for slices *i* = 1, 2.

We assign the first four unit vectors in the standard basis 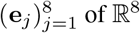 to cell type A, and the last four to cell type B. The top and bottom sides of *R*_i_(A) are assigned to features **e**_1_ and **e**_3_, while the left and right sides of the rectangle are assigned to features **e** and **e**. We make similar assignments for *R*_*i*_ (B) using the last four basis vectors. At each spot **s**, lying in some bounding rectangle *R* for its cell type, the feature assigned to **s** is a convex combination of the features decorating the top and bottom sides of *R*, plus the features decorating the left and right sides of *R*. In particular, for *R* = [*x*_min_, *x*_max_] × [*y*_min_, *y*_max_] and (*x, y*) ∈ *R*, the coefficients *λ*_x_, *λ*_y_ are defined as:

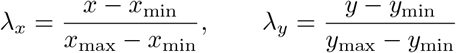

Thus, for a given cell type we have two gradients along the x and y direction which are scaled equivalently between the two slices **S**^(1)^ and **S**^(2)^ which we seek to align. The final feature vector within cell type A is therefore given as

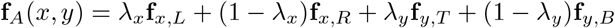

and repeat this for cell type B as

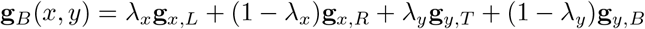

choosing features which are mutually-orthogonal–for simplicity 8-dimensional unit vectors **e**_1:8_. Between **S**^(1)^ and **S**^(2)^ these features are consistent, and *λ*_x_, *λ*_y_ ∈ [0, 1] gives the proportion of each feature depending on how far the (*x, y*) coordinate is along the axis of each ellipse within a given cell type. I.e. for (*x*_1_, *y*_1_) ∈ **S**^(1)^, (*x*_2_, *y*_2_) ∈ **S**^(2)^ we have that if 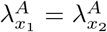 and 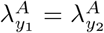 then **f**_A_(*x*_1_, *y*_1_) = **f**_A_(*x*_2_, *y*_2_) and the two features should be aligned between timepoints 1 and 2 (likewise for cell type B). As the features are chosen to be orthogonal, this alignment is unique. To model noise, we assume that the data we observe at each point (*x, y*) is distributed as 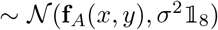 for cell type A and 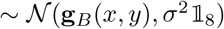 for cell type B for varying levels of *σ*.

## S8 Benchmarking methods

### PASTE [45]

By design, PASTE cannot align spatiotemporal data and does not infer cell growth and death because it uses balanced OT. For the purpose of benchmarking experiments in this work, we relax PASTE’s balanced constraints so that we optimize the unbalanced version of the PASTE objective. For the alignment matrix **Π** computed by relaxed PASTE, we then take the difference between the row sums of **Π** and the uniform distribution **g**_1_ (Eq. (4)) as PASTE’s inferred growth ***ξ***.

### moscot [21]

We use `moscot.problems.spatiotemporal.SpatioTemporalProblem` for solving spatiotemporal alignment problems in this work using moscot. For the alignment matrix **Π** computed by moscot, we take the difference between the row sums of **Π** and the uniform distribution **g**_1_ as moscot’s inferred growth ***ξ***. For benchmarking experiments related to growth *rates* instead of growth (§**??**), we use the numbers returned by `SpatioTemporalProblem.posterior_growth_rates` as moscot’s inferred growth *rates* over spots in timepoint *t*_1_.

### SLAT [44]

We use `scSLAT.model.run_SLAT` of the scSLAT package for computing an embedding for each spot in each timepoint. We then use `scSLAT.model.spatial_match` to compute an alignment between spots across the two timpoints. We construct a matrix **Π** such that **Π** _ij_^′^ = 1 if spot *i* in timepoint 1 is the best spot aligned to spot *j*^′^ in timepoint 2. We then divide **Π** by the number of spots at timpoint 1 to convert it into a semi-relaxed transport matrix. We take the difference between the row sums of **Π** and the uniform distribution **g**_1_ as SLAT’s inferred growth ***ξ***.

### STalign [8]

We first “rasterize” the spatial positions of our input data into two greyscale images. This effectively convolves a scatterplot of the spatial data with two-dimensional Gaussian noise, to produce an image approximating each tissue slice. We then ran STalign’s `LDDMM` on the pair of images, outputting a diffeomorphism *φ* (which maps the first slice to the second) and its inverse *φ*^−1^ (mapping the second slice to the first). We then use *φ*^−1^ to construct a transport plan **Π** as follows: for spot *j*^′^ in the second slice, we apply *φ*^−1^ to *s*_j_^′^, and select *s*_i_ in the first slice which is closest to *φ*^−1^(*s*_j_^′^). Then we set **Π** _ij_^′^ = 1, and repeat this procedure for each spot in the second slice, filling out each column of **Π** . The transport plan is thus a deterministic map from the spots of the second slice into those of the first. We then divide **Π** by the number of spots at timpoint 1 to convert it into a semi-relaxed transport matrix. We take the difference between the row sums of **Π** and the uniform distribution **g**_1_ as STalign’s inferred growth ***ξ***.

**Figure S1:**
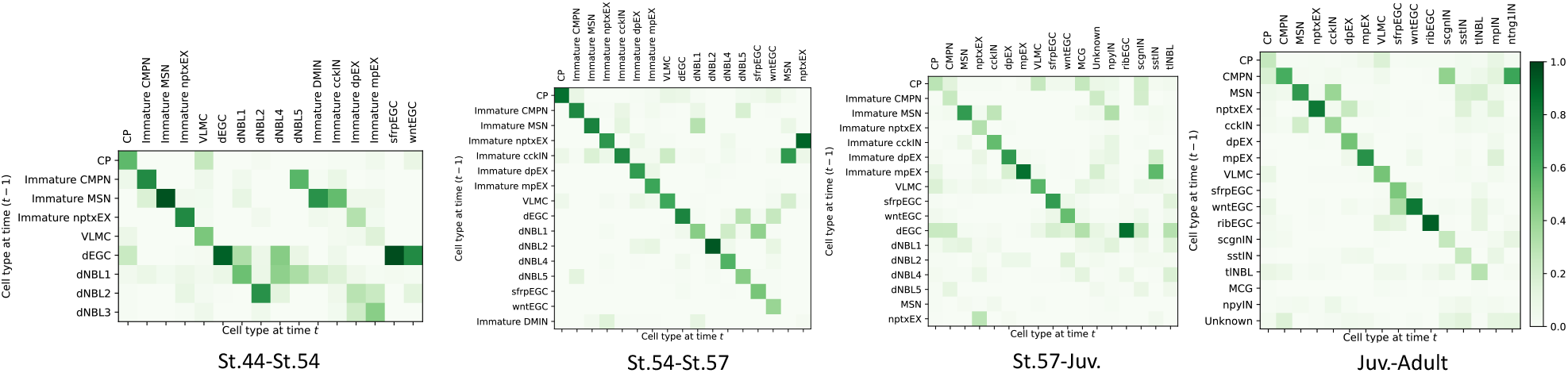
Cell type transition matrix at each pair of timepoints during axolotl brain development. Rows are cell types at the previous timepoint. Columns are cell types at the next timepoint. Matrices are column normalized.

## Footnotes

1 All matrices involved are subject to a consistent data-dependent normalization by matrix norms which ensure the terms of the gradient are scaled analogously, such that *α* = 0.5 roughly means both terms contribute equal weight for interpretability and robustness of the hyper-parameter defaults to different datasets.

